# Jagged-1^+^ Tregs Mediate Lymphatic Remodeling in Tumour-Draining Lymph Nodes

**DOI:** 10.1101/2024.06.10.598373

**Authors:** Jessie Z. Xu, Prudence PokWai Lui, Pei-Hsun Tsai, Hafsah Aziz, Boyu Xie, Inchul Cho, Manuela Terranova-Barberio, Chao Zheng Li, Katie Stoker, Hongqiang Yu, Alice E. Denton, Toby Lawrence, Sophia N. Karagiannis, Niwa Ali

## Abstract

The tumour-draining lymph nodes (tdLNs) are crucial sites for early immune surveillance of tumours. During tumour development, tdLNs undergo significant stromal remodelling that impacts lymph node (LN) architecture. The reconstruction of tdLNs is dependent on the crosstalk between regulatory T cells (Tregs) and resident stromal cells. The Notch ligand Jagged1 (Jag1), highly expressed in skin Tregs, is critical for facilitating stem cell-mediated tissue regeneration. However, the role of Jag1^pos^ Tregs in mediating the activity of tdLN-resident immune cells and stromal cells remains unexplored. Here, we used the B16-F10 tumour model to assess the role of Jag1^pos^ Tregs during tdLN development. During tumor progression, late-stage tdLN Tregs expressed higher levels of Jag1 than early-stage tdLNs. Conditional deletion of Jag1 in Tregs markedly restrained tdLN expansion, without influencing effector T cell abundance or activation profile. Transcriptomic analysis of tdLNs revealed downregulation of lymphatic endothelial cell (LEC)-related markers in mice with Treg-specific Jag1 ablation. Disruption of lymphatic networks in the tdLN was further confirmed by flow cytometric profiling and histological analysis. *In vivo* permeability assessment by FITC-dextran demonstrated perturbed lymphatic drainage, likely due to suppressed LN lymphangiogenesis. Finally, we identify the presence of Jag1^pos^ Tregs in tdLNs of human melanoma patients. Overall, our results highlight that crosstalk between Jag1^pos^ Tregs and LECs is required for lymphatic sinus expansion, which is critical for the formation of the tdLN niche during melanoma progression.

## INTRODUCTION

Regulatory T cells (Tregs) are key players in preventing inflammation and autoimmune disease, but their accumulation in the tumour microenvironment (TME) is highly correlated with poor prognosis in multiple cancer types.^1^ Beyond the primary tumour, tumour-draining lymph nodes (tdLNs) serve as key drainage sites essential for antigen recognition and distant metastasis.^2^ In both mice and humans, these sentinel tdLNs are highly enriched with Treg populations exhibiting a unique transcriptomic profile compared to paired non-tdLNs.^3,4^ This enrichment poses clinical opportunities for local tdLN immunotherapies, an alternative to targeting tumours with a higher immunosuppressive status.^5^

Tumour development is accompanied by tdLN enlargement and lymphatic remodelling to favour a pre-metastatic lymphovascular niche conducive to tumour cell colonization.^6^ Lymphatic vessels are vital for maintaining the structural and functional stability of the lymph node in an organotypic manner.^7^ Mounting evidence points towards a major role for Tregs in governing lymphovascular function and cellular homeostasis. Tregs can directly engage with LT beta receptor (LTβR) on lymphatic endothelial cells (LECs) to mediate leukocyte transendothelial migration into draining LNs.^8–10^ Another study demonstrated that engagement between PD-1 in Tregs with PD-L1 expressing LECs can facilitate Treg migration.^11^ Additionally, Tregs are endowed with the capacity to reverse the pathological hallmarks of lymphedema, including impaired lymphatic transport capacity.^12^ Notably, short-term depletion of tumour infiltrating Tregs preferentially impacts the transcriptome of fibroblasts and endothelial cells in the TME, rather than lymphoid and myeloid immune effector cell subsets. These findings underscore the strong Treg connectivity to non-immune accessory cell types during the evolution of an anti-tumour immune response.^13^

In recent years, it has become evident that Tregs residing outside secondary lymphoid organs possess distinct tissue-specific functions. Previously, we identified the Notch ligand, Jagged1 (Jag1), as a preferentially expressed receptor on skin Tregs that facilitates epithelial stem cell mediated tissue regeneration.^14^ The Notch pathway is a compelling candidate for mediating Treg function in tdLNs, given the intimate association with thymic Treg generation and the control of Treg suppressive capacity in the periphery.^15^ Furthermore, Jag1 expression outside of T cells, particularly in blood vessels, acts as a potent proangiogenic regulator.^16,17^ Whether this pathway is also utilized by LN-residing Tregs in the context of tumorigenesis or the maintenance of stromal cell dynamics in tdLNs has not been explored.

Here, we show that during murine melanoma progression, Jag1^pos^ Tregs are highly activated and accumulate in late-stage tdLNs, but not in non-tdLNs. Cell specific ablation of Jag1 in Tregs (Foxp3^ΔJag1^) results in a failure of tdLN expansion, accompanied by a global reduction in LN cellularity and impaired lymphatic drainage. Transcriptomic and cellular analysis of Foxp3^ΔJag1^ and Foxp3^Ctrl^ tdLNs revealed dysregulated expression of LEC-associated markers. We also report selective expression of Jag1 on Tregs in human sentinel lymph nodes from melanoma patients. Overall, our findings reveal that Jag1^pos^ Tregs in the tdLN are essential for maintaining lymphatic endothelial cell homeostasis during melanoma tumour development.

## RESULTS

### Jag1^pos^ Tregs accumulate in late tumour-draining lymph nodes

Previously, we have described a subpopulation of Tregs resident in skin that express the Notch ligand, Jagged-1 (Jag1).^14^ To explore if Jag1-expressing Tregs are present in the tumour microenvironment (TME) and/or exhibit any tumour-associated changes, we utilised the *in vivo* B16-F10 melanoma model. Two time points were selected for flow cytometric phenotyping of subcutaneously implanted tumours in C57BL/6J mice. The ‘Early’ time point, at day 10-11 post-tumour implantation was identified when animals developed minimally palpable tumours, whereas ‘Late’ stage was defined when tumour volumes exceeded 200m^3^, generally on day 14-17 (**Figure 1A**). In the corresponding tdLNs of tumour implanted animals, Late stage Tregs expressed higher Jag1 levels than Early stage Tregs (**Figure 1B-C**; full gating strategy is shown in **Figure S1**). This increase was reflected in an overall higher abundance of Jag1^pos^ Tregs in Late tdLNs (**Figure 1C**). To assess the phenotypic similarity between Jag1^pos^ Tregs and the remaining Jag1^neg^ Treg fraction, we profiled the expression of activation-associated phenotypic markers CTLA4, ICOS, CD25, and Ki67. Intriguingly, Jag1^pos^ Tregs in tdLNs exhibited higher levels of Ki67 than Jag1^neg^ Tregs at both timepoints (**Figure 1D**). This was also true for other markers, which were upregulated in the Jag1^pos^ fraction.

**Figure 1.**
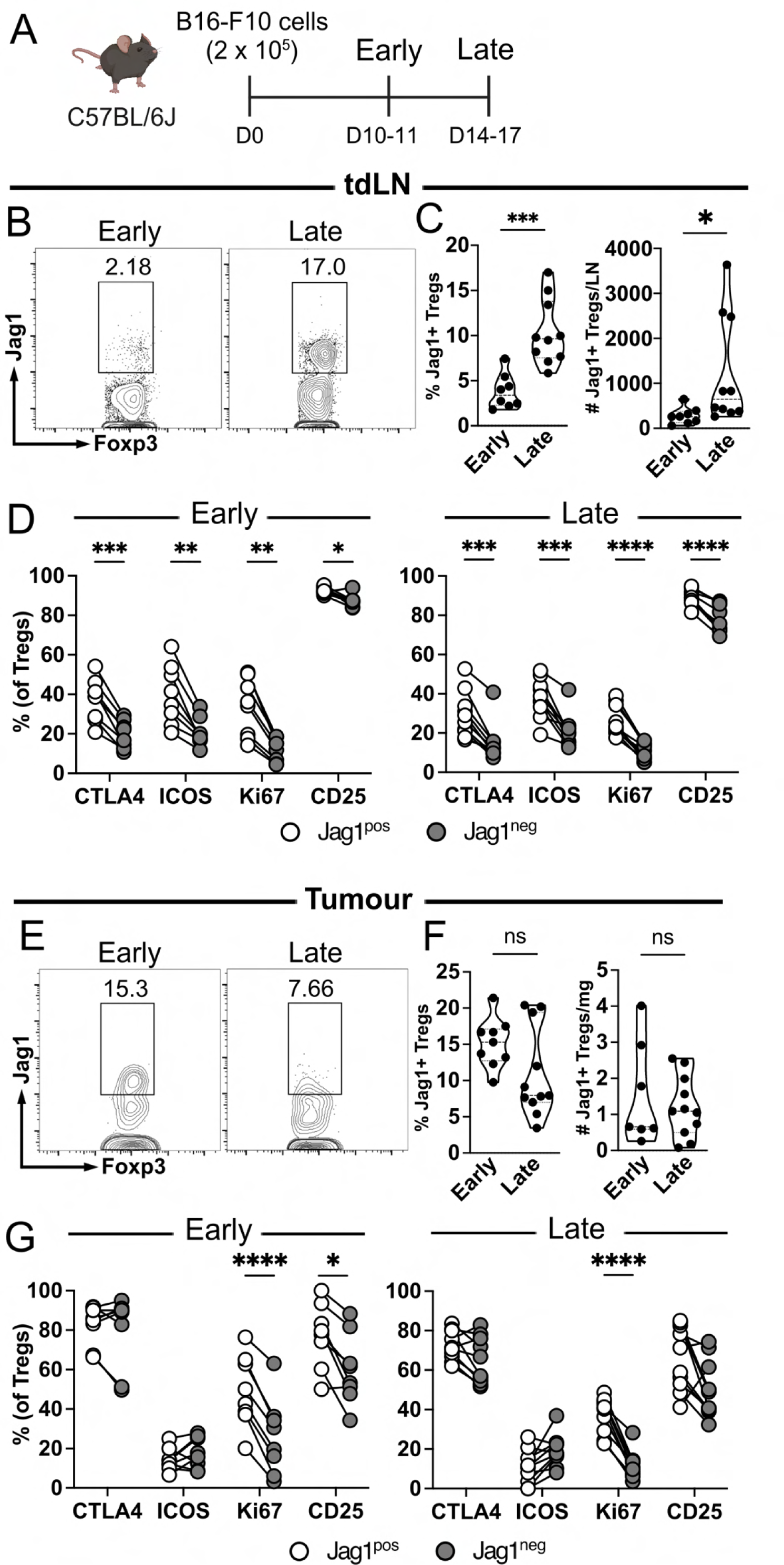
Jag1^pos^ Tregs accumulate in late tumour-draining lymph nodes. **A)** Schematic of experimental timeline. Wildtype mice were intradermally injected with B16F10 melanoma cells on day 0 (D0) and harvested at early (D10-11) or late (D14-17) time points. **B)** Representative flow plots and **C)** quantification of % Jag1 expression on Tregs and absolute number of Jag1^pos^ Tregs per tdLN between early and late time points. **D)** % expression of activation markers (CTLA4, ICOS, Ki67, CD25) between Jag1^pos^ and Jag1^neg^ Tregs in early and late tdLN. **E)** Representative flow plots and **F)** quantification of % Jag1 expression on Tregs and absolute number of Jag1^pos^ Tregs per mg of tumour at early and late stage. **G)** % expression of activation markers (CTLA4, ICOS, Ki67, CD25) between Jag1^pos^ and Jag1^neg^ Tregs in early and late tumour. Data are representative of two independent experiments. Results were presented as mean ± SEM. Statistics were calculated by unpaired t-test for (C, F), and by multiple paired t-test for (D, G). ****p<0.0001, ***p<0.001, **p<0.01, *p<0.05, ns p>0.05.

We further assessed the same parameters in primary tumours. In contrast to the tdLN site, early and Late tumour-infiltrating Tregs expressed appreciable levels of Jag1 (**Figure 1E-F**). However, there was no significant change in their proportion within the Foxp3+ Treg fraction, nor in their overall abundance between these time points (**Figure 1F).** Similarly to the tdLNs, both Early and Late Jag1^pos^ Tregs displayed higher levels of the proliferation marker Ki67, relative to their Jag1^neg^ counterparts (**Figure 1G**). Late Jag1^pos^ Tregs also expressed higher levels of CD25, suggesting that tumour infiltrating Jag1^pos^ Tregs may represent a highly activated subpopulation. Overall, these findings suggest that Jag1 may be a putative marker to delineate the activation status of Tregs.

Taken together, these findings reveal that Jag1-expressing Tregs are present amongst the tumour infiltrating immune compartment but are particularly prominent within draining lymph nodes. Further characterisation highlighted Jag1^pos^ Tregs as a highly activated subpopulation and identifies the Late stage of tumour development as a timeframe where these cells accumulate in the tdLNs.

### Conditional deletion of Jag1 in Tregs restrains lymph node expansion

The accumulation of highly activated Jag1^pos^ Tregs in Late tdLNs raised the notion of whether these cells play a role in limiting tumour immune surveillance and/or impacting melanoma progression. To study the function of Jag1^pos^ Tregs on melanoma progression, we utilized an inducible Foxp3-driven Cre-ert2 Jag1-LoxP system whereby the Jag1 allele is deleted in a lineage-specific and temporal fashion in Foxp3-expressing Tregs. To induce Jag1 deletion, we administered tamoxifen (TAM) intraperitoneally for 5 consecutive days and then rested the animals for a 9 day washout period (**Figure 2A).**^18,19^ Subsequently, B16F10 melanoma cells were implanted in Foxp3^CreErt2^Jag1^fl/fl^ and control Foxp3^CreErt2^Jag1^fl/wt^ mice, hereafter referred to as Foxp3^ΔJag1^ and Foxp3^Ctrl^, respectively. Under these conditions, Jag1^pos^ Tregs had no major impact on overall tumour kinetics given the unchanged tumour volume relative to controls (**Figure 2B-C).** At endpoint, tumour weight also remained unchanged (**Figure 2C**), indicating that Treg-specific loss of Jag1 did not impact tumour growth. Additionally, the total cellularity and abundance of major cell subpopulations infiltrating the tumour were unaffected (**Figure 2D-E).**

**Figure 2.**
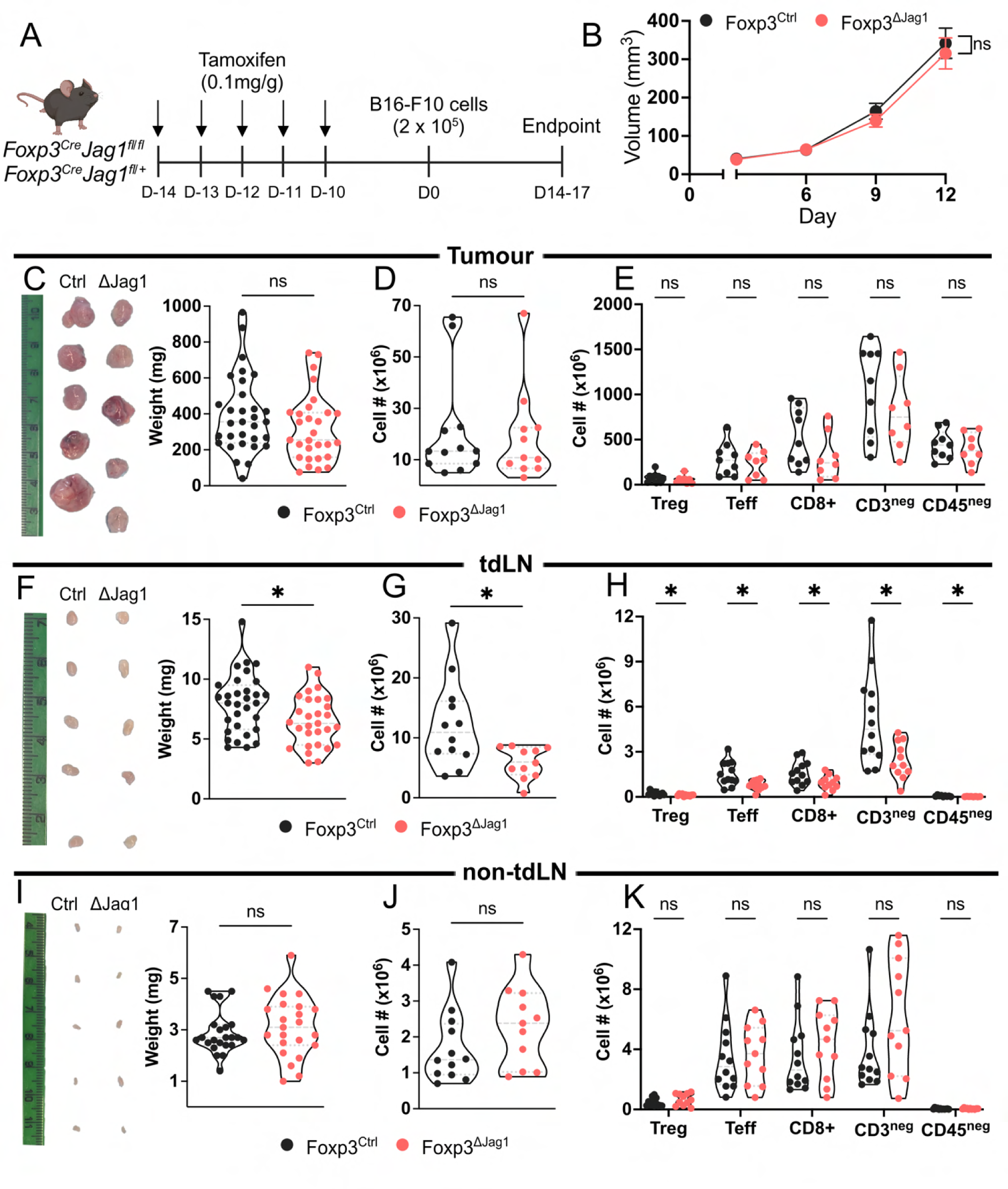
Conditional deletion of Jag1 in Tregs restrains lymph node expansion. **A)** Schematic of experimental timeline. Foxp3^ΔJag1^ and Foxp3^Ctrl^ mice received five consecutive doses of tamoxifen (0.1mg/g) between Day −14 and Day −10, then were injected with B16F10 cells on D0 before being harvested on D14-17. **B)** Tumour volume (mm^3^) between Foxp3^ΔJag1^ and Foxp3^Ctrl^ groups as calculated by (1/2)(length*width^2^) on D3, D6, D9 and D12 post tumour inoculation. **C)** Representative images and weight (mg) of Foxp3^ΔJag1^ and Foxp3^Ctrl^ tumours at endpoint. **D-E)** Absolute number of **D)** total cells and **E)** Treg, Teff, CD8+ T cell, CD3- and CD45-subpopulations between Foxp3^ΔJag1^ and Foxp3^Ctrl^ tumours at endpoint. **F)** Representative images and weight (mg) of Foxp3^ΔJag1^ and Foxp3^Ctrl^ tdLNs at endpoint. **G-H)** Absolute number of **G)** total cells and **H)** Treg, Teff, CD8+ T cell, CD3- and CD45-subpopulations between Foxp3^ΔJag1^ and Foxp3^Ctrl^ tdLNs at endpoint. **I)** Representative images and weight (mg) of Foxp3^ΔJag1^ and Foxp3^Ctrl^ non-tdLNs at endpoint. **J-K)** Absolute number of **J)** total cells and **K)** Treg, Teff, CD8+ T cell, CD3- and CD45-subpopulations between Foxp3^ΔJag1^ and Foxp3^Ctrl^ non-tdLNs at endpoint. Data are representative of three independent experiments. Results were presented as mean ± SEM. Statistics were calculated by unpaired t-test for (C-D, F-G, I-J), by two-way ANOVA for (B), and by multiple unpaired t-test for (E, H, K). *p<0.05, ns p>0.05.

In the B16F10 model, tumour growth is accompanied by increasing tdLN weight and enlargement of lymphatic vessels.^6^ Therefore, we wondered if tumour-involved tdLNs were also affected by Jag1 deletion in Tregs. Interestingly, we observed significantly reduced tdLN mass in Foxp3^ΔJag1^ relative to Foxp3^Ctrl^ controls (**Figure 2F**). The reduced mass of tdLNs was also accompanied by a reduction in total cellularity in Foxp3^ΔJag1^ animals (**Figure 2G**). Upon further characterisation of tdLNs, we found a largely uniform reduction in the total cellularity of multiple immune and stromal cell subpopulations (**Figure 2H)**. Conversely, the weight and cellularity differences were absent in non-tdLNs (**Figure 2I-K**), indicating the impact on lymph node expansion is dependent on the presence of a proximal tumour. To confirm this, we performed full thickness skin wounding in non-tumour implanted Foxp3^ΔJag1^ and Foxp3^Ctrl^ animals to induce a fulminant inflammatory response. Indeed, assessment of wound-draining LNs (wound-dLNs) displayed no difference in total cellularity or the abundance of major cellular fractions (**Figure S2A-B**), suggesting Jag1^pos^ Treg-mediated expansion of lymph nodes is confined to the tumour context. Collectively, we show that Jag1^pos^ Tregs are required for the maintenance of tdLN size and overall cellularity during late stage melanoma tumour progression.

### Loss of Jag1 in Tregs has no major impact on T cell-mediated immunity

To determine whether the failure of tdLN expansion in Foxp3^ΔJag1^ mice is related to major hallmarks of disruption in T cell-mediated immunity, we performed flow cytometric profiling in tumour and LN tissues. Tumour-infiltrating Treg numbers were comparable between conditions, as were Teff and CD8+ T cells numbers (**Figure 3A-B**). Accordingly, Teff:Treg and CD8:Treg ratios remained unchanged (**Figure 3C**), suggesting that the overall balance between pro- and anti-tumour immunogenicity was maintained. Given Jag1^pos^ Tregs adopt a highly activated profile in the context of tumour progression in wild type mice (**Figure 1**), we assessed the expression of activation markers in Tregs, Teffs and CD8+ T cells between endpoint Foxp3^ΔJag1^ and Foxp3^Ctrl^ tissues. We observed no differences in all activation markers between Tregs, Teffs, and CD8+ T cells infiltrating tumours (**Figure 3D-F**). This was also true for protein expression on a per cell basis, as measured by the mean fluorescence intensity (MFI) (**Figure S3A-C**).

**Figure 3.**
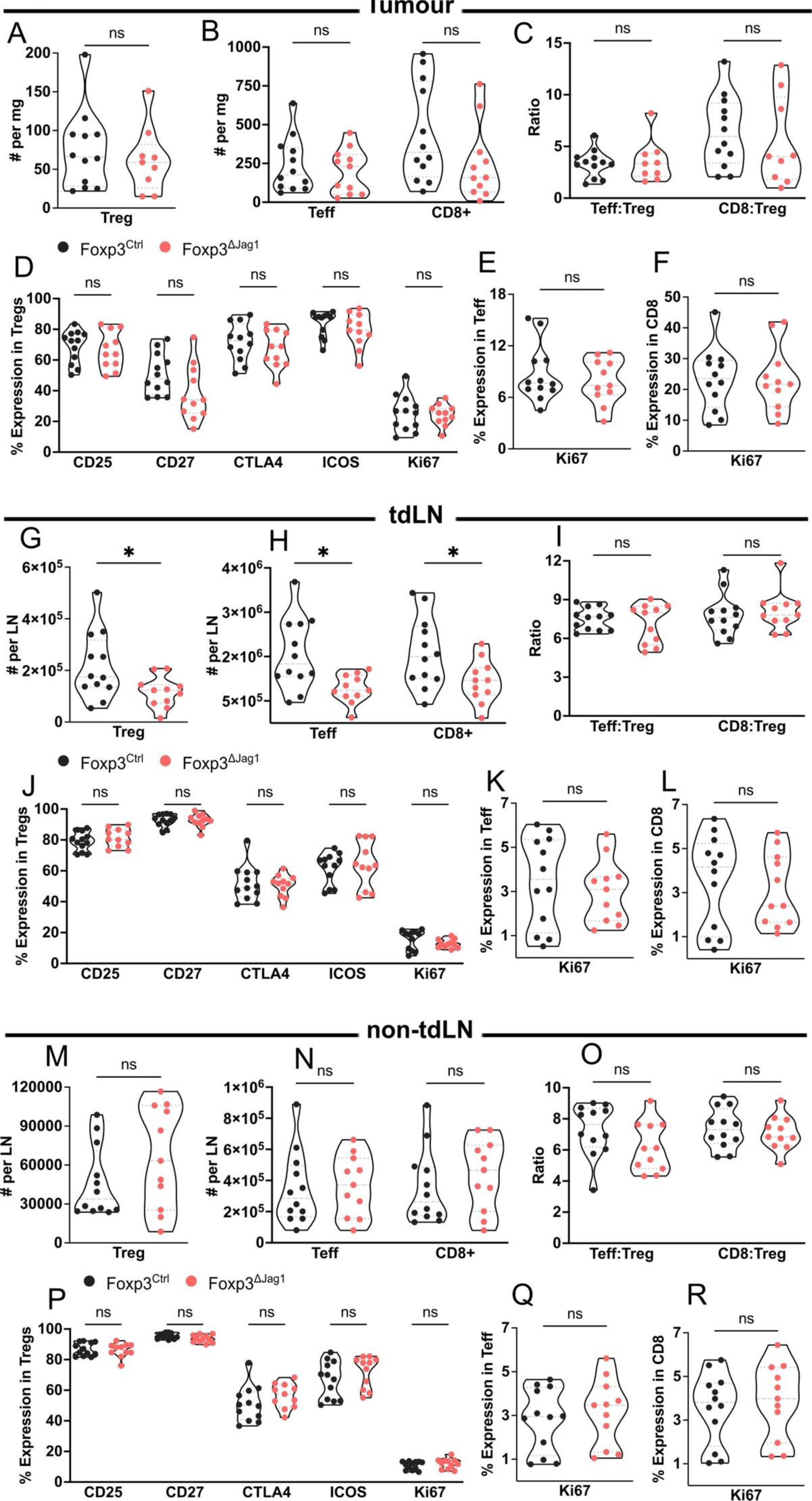
Loss of Jag1 in Tregs has no major impact on T cell-mediated immunity. **A-B)** Absolute number quantification of **A)** Tregs and **B)** Teffs and CD8+ T cells in endpoint Foxp3^ΔJag1^ and Foxp3^Ctrl^ tumour digests. **C)** Teff:Treg and CD8:Treg ratios between endpoint Foxp3^ΔJag1^ and Foxp3^Ctrl^ tumour. **D)** Flow cytometric quantification of % expression of activation markers CD25, CD27, CTLA4, ICOS and KI67 on endpoint Foxp3^ΔJag1^ and Foxp3^Ctrl^ tumour Tregs. **E-F)** % Ki67 expression on endpoint Foxp3^ΔJag1^ and Foxp3^Ctrl^ tumour **E)** Teffs and **F)** CD8+ T cells. **G-H)** Absolute number quantification of **G)** Tregs and **H)** Teffs and CD8+ T cells in endpoint Foxp3^ΔJag1^ and Foxp3^Ctrl^ tdLNs. **I)** Teff:Treg and CD8:Treg ratios between endpoint Foxp3^ΔJag1^ and Foxp3^Ctrl^ tdLNs. **J)** Flow cytometric quantification of % expression of activation markers CD25, CD27, CTLA4, ICOS and KI67 on endpoint Foxp3^ΔJag1^ and Foxp3^Ctrl^ tdLN Tregs. **K-L)** % Ki67 expression on endpoint Foxp3^ΔJag1^ and Foxp3^Ctrl^ tdLN **K)** Teffs and **L)** CD8+ T cells. **M-N)** Absolute number quantification of **M)** Tregs and **N)** Teffs and CD8+ T cells in endpoint Foxp3^ΔJag1^ and Foxp3^Ctrl^ non-tdLNs. **O)** Teff:Treg and CD8:Treg ratios between endpoint Foxp3^ΔJag1^ and Foxp3^Ctrl^ non-tdLNs. **P)** Flow cytometric quantification of % expression of activation markers CD25, CD27, CTLA4, ICOS and KI67 on endpoint Foxp3^ΔJag1^ and Foxp3^Ctrl^ non-tdLN Tregs. **Q-R)** % Ki67 expression on endpoint Foxp3^ΔJag1^ and Foxp3^Ctrl^ non-tdLN **Q)** Teffs and **R)** CD8+ T cells. Data are representative of three independent experiments. Results were presented as mean ± SEM. Statistics were calculated by unpaired t-test for (A, E, F, G, K, L, M, Q, and R), and by multiple unpaired t-test for (B-D, H-J, and N-P). *p<0.05, ns p>0.05.

By contrast, in Foxp3^ΔJag1^ tdLNs we noted a reduction in the abundance of Tregs, accompanied by similarly reduced numbers of Teffs and CD8s (**Figure 3G-H**). Nonetheless, this reciprocal reduction did not impact Teff:Treg and CD8:Treg ratios (**Figure 3I**). Similarly, for tdLNs of Foxp3^ΔJag1^ and Foxp3^Ctrl^ animals, both the proportion and MFI of activation marker expression were unchanged (**Figure 3J-L, and S3D-F**). In addition, the abundance of T cell subpopulations and the ratios of Teff:Treg and CD8:Treg were unaffected in non-tdLNs (**Figure 3M-O**). The endpoint non-tdLN compartment also showed comparable expression of activation markers both by proportion (**Figure 3P-R)** and MFI **(Figure S3G-I).**

Given the dynamic cellular nature of tdLNs, we also characterized an earlier time point, day 4, to assess any temporal changes of T cell subpopulations. Unlike endpoint tdLNs, day 4 tdLNs did not differ in Treg and Teff numbers in Foxp3^ΔJag1^ and Foxp3^Ctrl^ animals (**Figure S3J**). As a result, the Teff:Treg ratio was also consistent between the two conditions (**Figure S3K)**. Interestingly, day 4 tdLN Tregs displayed elevated levels of Ki67 (**Figure S3L)**, which corresponded to a similar change in MFI (**Figure S3M)**. This proliferative phenotype was not observed in tdLN Teffs based on the proportion and MFI of Ki67 expression (**Figure S3N-O)**.

Overall, our findings suggest that loss of Jag1 in Tregs has no major impact on the activation status of T cell subsets implicated in regulating the anti-tumour immune response. Thus, we ruled out that Jag1^pos^ Tregs play a major role in regulating the proliferation and activation of effector immune cells in tdLNs.

### Lymph node Jag1^pos^ Treg promote stromal-related gene signatures

To identify the key cell types and/or signaling pathways involved in Jag1^pos^ Treg mediated expansion of tdLNs, we performed bulk RNA-sequencing of Foxp3^ΔJag1^ and Foxp3^Ctrl^ tdLN and tumour tissues. Given that Jag1 is a prominent ligand of the Notch signaling pathway, we first assessed if Notch related genes were impacted. In tdLNs, we identified 653 differentially expressed genes (DEGs) with p-value <0.05. Of these, 33 DEGs (∼5.3%) were shared with a curated Notch-target gene set (p-value of the overlap = 4.94 × 10^-6^) (**Figure 4A**), indicating that deletion of Jag1 in Tregs led to aberrant Notch signaling within the tdLN environment.

**Figure 4.**
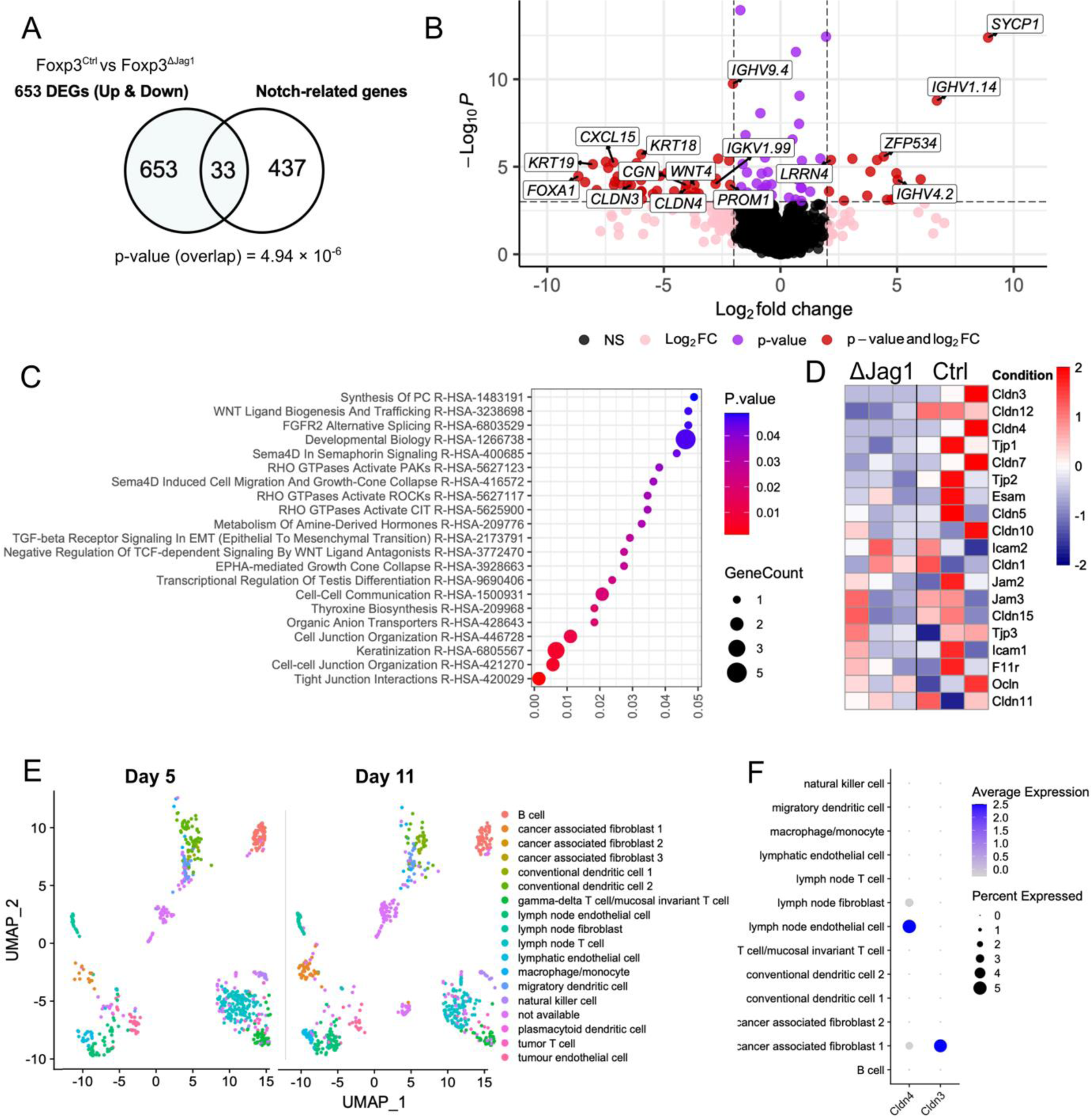
Lymph node Jag1^pos^ Treg promote stromal-related gene signatures. **A)** Venn diagram showing the overlap between tdLN bulk RNA-Seq total DEGs (p-value <0.05) and known Notch-related genes. P-value represents the significance of the overlap as calculated by chi-squared test. **B)** Differential gene analysis between Foxp3^ΔJag1^ and Foxp3^Ctrl^ tdLNs. A negative fold change denotes Foxp3^Ctrl^-upregulated genes and a positive fold change denotes Foxp3^Ctrl^-downregulated genes. Dots in red are individual DEGs with >2 log2 fold change and <0.05 p-adjusted value. **C)** Reactome pathway enrichment analysis using tdLN Foxp3^Ctrl^-upregulated DEGs (p-adjusted value <0.05). **D)** Heatmap comparing expression of tight junction family genes between Foxp3^ΔJag1^ (ΔJag1) and Foxp3^Ctrl^ (Ctrl) tdLNs. **E)** UMAP showing temporal changes in immune and stromal cell composition between D5 and D11 tdLN. F). Dot plot showing cell type-specific expression of Jag1^pos^ Treg-associated DEGs identified from tdLN bulk RNA-Seq in previously published scRNA-Seq dataset of B16F10 tdLN.

Differential analyses between Foxp3^ΔJag1^ vs Foxp3^Ctrl^ tdLNs revealed an increased abundance of genes related to cell junction organization (*CGN*, *CLDN3*, *CLDN4*, *CDH1, TJP1, DSP*) and epithelial to mesenchymal (EMT) transition in cancer (*KRT8, KRT18*, *KRT19*) (**Figure 4B**). In addition, we detected fluctuating expression in several immunoglobulin gene segments (*IGHV9.4, IGKV1.99,IGHV4.2, IGHV1.14)* (**Figure 4B)**, related to B cell receptor re-arrangement. Reactome enrichment analysis enriched terms such as “tight junction interactions” and “cell-cell junction organization” associated with upregulated *CLDN3* and *CLDN4* in Foxp3^Ctrl^ tdLNs (**Figure 4C)**. *CLDN3* and *CLDN4* are cell adhesion genes that encode for tight junction proteins between epithelial or endothelial cell sheets to reinforce barrier function.^20^ We also noted other key tight junction transcripts were over-represented in Foxp3^Ctrl^ tdLNs, such as *CLDN12, TJP1, CLDN7,* and *TJP3* **(Figure 4D)**. Reactome pathway analysis also identified Wnt/β-catenin as an enriched pathway in Foxp3^Ctrl^ tdLNs (**Figure 4C**), a pathway implicated in both lymphangiogenesis and lymphatic vascular development.^21,22^ These results suggest a possible interaction between Jag1^pos^ Treg and stromal cells via the Wnt/β-catenin pathway in the tdLN.

To infer cell-type specificity of claudin transcripts in the lymph node, we leveraged a publicly available scRNA-Seq dataset of CD45+ immune cells and CD45-stromal cells isolated from day 5 and 11 of tdLNs from B16-F10 implanted mice. To reconstruct the cellular composition of a developing tdLN, we plotted UMAPs showing clustering of cell types in day 5, 8 and 11. In particular, the CD45-CD31+ endothelial population were enriched in tdLNs at endpoint (**Figure 4E**). As expected, we found that the *CLDN4* and *CLDN3* were almost exclusively expressed in lymphatic endothelial cells (LECs) and cancer-associated fibroblasts, respectively (**Figure 4F)**. Prior studies have indicated the global upregulation of these adhesion-related genes can diminish tdLN permeability and mediate LEC adhesion to the extracellular matrix.^6^ As such, our results highlight that Jag1-expressing Tregs may modulate the function and properties of tumour-associated stromal cell subsets.

Conversely, the number of DEGs discovered in the tumour showed insignificant overlap with known Notch-target transcripts (p-value of the overlap = 0.668) (**Figure S4A**). Upon filtering out genes with adjusted p-value (P_adj_) > 0.05, the remaining DEGs were predominantly downregulated in Foxp3^ΔJag1^ tumours and associated with the regulation of adipogenesis (*ADIPOQ, LEP, PLIN1)* (**Figure S4B**). Further enrichment analysis also highlighted Reactome pathways related to lipid metabolism (**Figure S4C**), suggesting that Jag1^pos^ Tregs may modulate cancer-associated adipocytes in the TME.

Taken together, our transcriptomic analysis between Foxp3^ΔJag1^ and Foxp3^Ctrl^ conditions identified a significantly greater magnitude of DEGs and dysregulated Notch signalling in tdLNs relative to tumours. This is also consistent with our *in vivo* functional data where we note a highly significant impact on tdLN expansion. Given these findings, we focused our next series of experiments on exploring if lymph node stromal cell dynamics are under the control of Jag1^pos^ Tregs during tumour progression.

### Jag1^pos^ Tregs are required for tdLN lymphangiogenesis

To validate our *in silico* findings on LN stromal cell subtypes, we performed flow cytometric profiling of the CD45 negative fraction of Foxp3^ΔJag1^ and Foxp3^Ctrl^ tdLNs on both day 4 and at tumour endpoint. Day 4 was chosen as the earlier time point as it corresponds to the beginning of LN enlargement when stromal cell populations are most proliferative.^6^ We then utilised well established protocols and cell surface expressed markers to delineate stromal cell subsets, namely, LECs as CD31+PDPN+ double positive, blood endothelial cells (BECs) as CD31+ single positive, and fibroblastic reticular cells (FRCs) as PDPN+ single positive.^10,23,24^ In line with our RNA-Seq data, the proportion of LECs was significantly reduced in endpoint Foxp3^ΔJag1^ tdLNs, while FRCs and BECs were unaffected (**Figure 5A-B**). Meanwhile, the proportions of LEC, FRC and BEC were all similar between the two conditions on Day 4 (**Figure S5A).**

**Figure 5.**
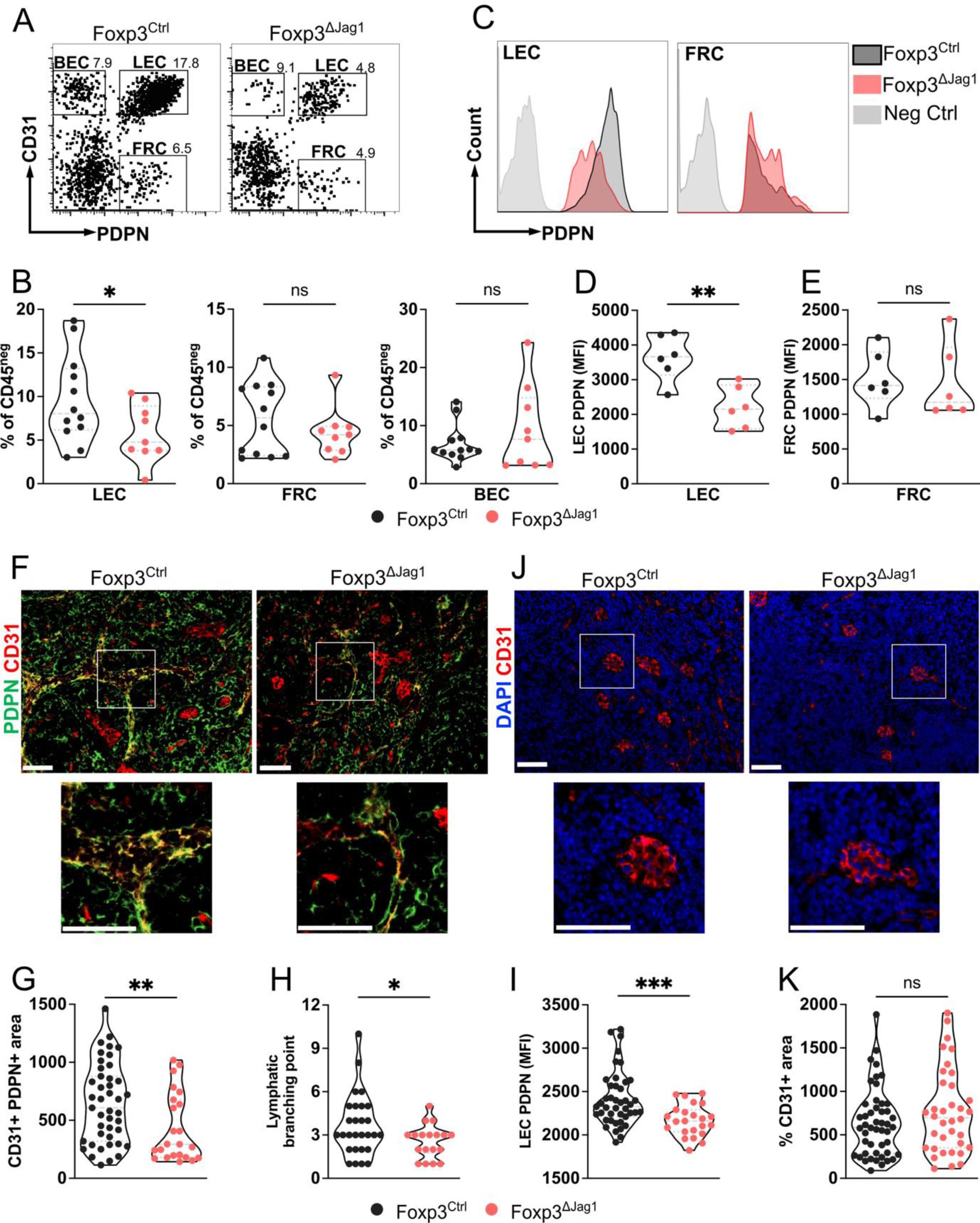
Jag1^pos^ Treg are required for tdLN lymphangiogenesis. **A)** Representative flow plots and **B)** flow cytometric quantification of the proportion of major CD45-stromal subpopulations (LECs, FRCs, BECs) out of total CD45-cells between endpoint Foxp3^ΔJag1^ and Foxp3^Ctrl^ tdLN. **C)** Representative flow plot and quantification of MFI PDPN on **D)** LECs and **E)** FRCs in Foxp3^ΔJag1^ and Foxp3^Ctrl^ tdLNs. **F)** Immunofluorescence staining of PDPN/CD31 double positive lymphatic vessels in Foxp3^ΔJag1^ and Foxp3^Ctrl^ tdLN at endpoint. **F)** *In situ* quantification of lymphatic vessel area, **H)** number of branching point and **I)** MFI PDPN per ROI of tdLN. 50 µm scale bar. **J)** Immunofluorescence staining and **K)** quantification of CD31+ blood vessels per tdLN region of interest. 50 µm scale bar. Data are representative of 2-3 independent experiments. Statistics were calculated by unpaired t-test. ***p<0.001, **p<0.01, *p<0.05, ns p>0.05.

We next sought to assess the per cell expression level of the PDPN protein in LECs (**Figure 5C**). This measure provides an indication of LEC activation and their concurrent capacity to maintain lymphatic vascular homeostasis.^25^ Previous studies have also shown a correlation between increased PDPN expression and LEC expansion in the LN.^26,27^ Strikingly, the MFI of PDPN was drastically reduced in LECs, but not FRCs, analysed from endpoint Foxp3^ΔJag1^ tdLNs, relative to Foxp3^Ctrl^ controls (**Figure 5D-E**). The reduction in PDPN MFI, however, was not present in LECs isolated from Day 4 Foxp3^ΔJag1^ tdLNs **(Figure S5B**). Similar to endpoint, FRCs showed no change in the MFI of PDPN on upon loss of Jag1 in Tregs (**Figure S5C**). This observation is highly suggestive of disrupted lymphatic vessel integrity in the absence of Jag1^pos^ Tregs.

We next quantified CD31+PDPN+ cells in endpoint tdLNs by immunofluorescence to measure the total LEC area occupied *in situ.* In line with our flow cytometric observation, LEC numbers were significantly reduced in Foxp3^ΔJag1^ tdLNs (**Figure 5F-G**). This was further confirmed by assessment of LYVE-1 (lymphatic vessel endothelial receptor 1) expression (**Figure S5D**), a specific marker for lymphatic endothelium.^28^ Our analyses revealed both the proportion of LYVE-1+ occupied regions and total vessel length were markedly reduced in Foxp3^ΔJag1^ tdLNs (**Figure S5E-F**). There were also considerably more branching of lymphatic vessels in Foxp3^Ctrl^ controls compared to Foxp3^ΔJag1^ tdLNs, a measure of lymphangiogenesis (**Figure 5H**). In addition, the observation of PDPN MFI reduction in Foxp3^ΔJag1^ tdLNs was also reflected when quantified by immunofluorescence (**Figure 5I**). Meanwhile, enumeration of CD31-expressing BECs in tdLNs revealed no change in the total area occupied between conditions (**Figure 5J-K**). Overall, our findings demonstrate that Jag1-expressing Tregs preferentially modulate LEC dynamics and consequent lymphangiogenesis in tdLNs.

### Jag1+ Tregs modulate lymphatic drainage to tdLNs

Thus far, we have identified a role for Jag1^pos^ Tregs in remodelling lymphatic networks in tdLNs during the late stages of tumour progression. To evaluate whether lymphatic vessel drainage and permeability was impaired upon Jag1 deletion in Tregs, we utilised a fluorescent tracing approach with FITC dextran.^7,29–31^ We administered 4kDa and 70kDa FITC-labelled dextrans *in vivo* into the footpad of endpoint tumour-bearing mice, 10 minutes before harvest (**Figure S6A**). Fluorescence quantification revealed an increasing trend of 4kDa flux in Foxp3^ΔJag1^ tdLN relative to controls (**Figure S6B**). Meanwhile, the normalised flux rate of 70kDa dextran showed a decreasing trend in Foxp3^ΔJag1^ tdLN compared to control (**Figure S6C**), indicating that passage of larger molecules and infiltrating cells may be largely interrupted. Indeed, previous studies have also demonstrated lymphatic drainage is highly dependent on LN lymphangiogenesis.^32^ Given that high lymphatic drainage positively correlates with cellular infiltration,^7^ this may have contributed to the lack of tissue expansion seen in Foxp3^ΔJag1^ tdLNs (**Figure 2F-H**). As such, our findings suggest that Jag1^pos^ Tregs may functionally modulate lymphatic drainage to tdLNs, likely via promotion of lymphangiogenesis.

### High Jag1 score on Tregs correlates with worse patient prognosis

We next considered the relevance of our murine data to human cancer, particularly in melanoma. We sought to address whether the Treg Jag1-mediated pathway is a reliable indicator of survival in melanoma patients. To this end, we used human orthologs of DEGs from our murine tdLN RNA-Seq to derive a Treg-Jag1 associated transcriptome profile to create a Jag1 scoring system (**Figure 6A**). We then scored TCGA cancer patient transcriptome data based on enrichment of Jag1-related profiles. Of the 350 melanoma patients, 117 stratified into Low Jag1 score, and 115 as High Jag1 scoring. From this TCGA cohort, patients in the Low Jag1 subgroup had longer overall survival times than those in the High Jag1 subgroup, according to the Kaplan-Meier analysis (p = 0.0048; **Figure 6B**). Of note, poor survival associated with Jag1^pos^ Tregs was also present in breast cancer (p = 0.048; **Figure 6C**), but no correlation was observed in ovarian and colorectal cancer (p = 0.77 and p = 0.67, respectively; **Figure 6D-E**).

**Figure 6.**
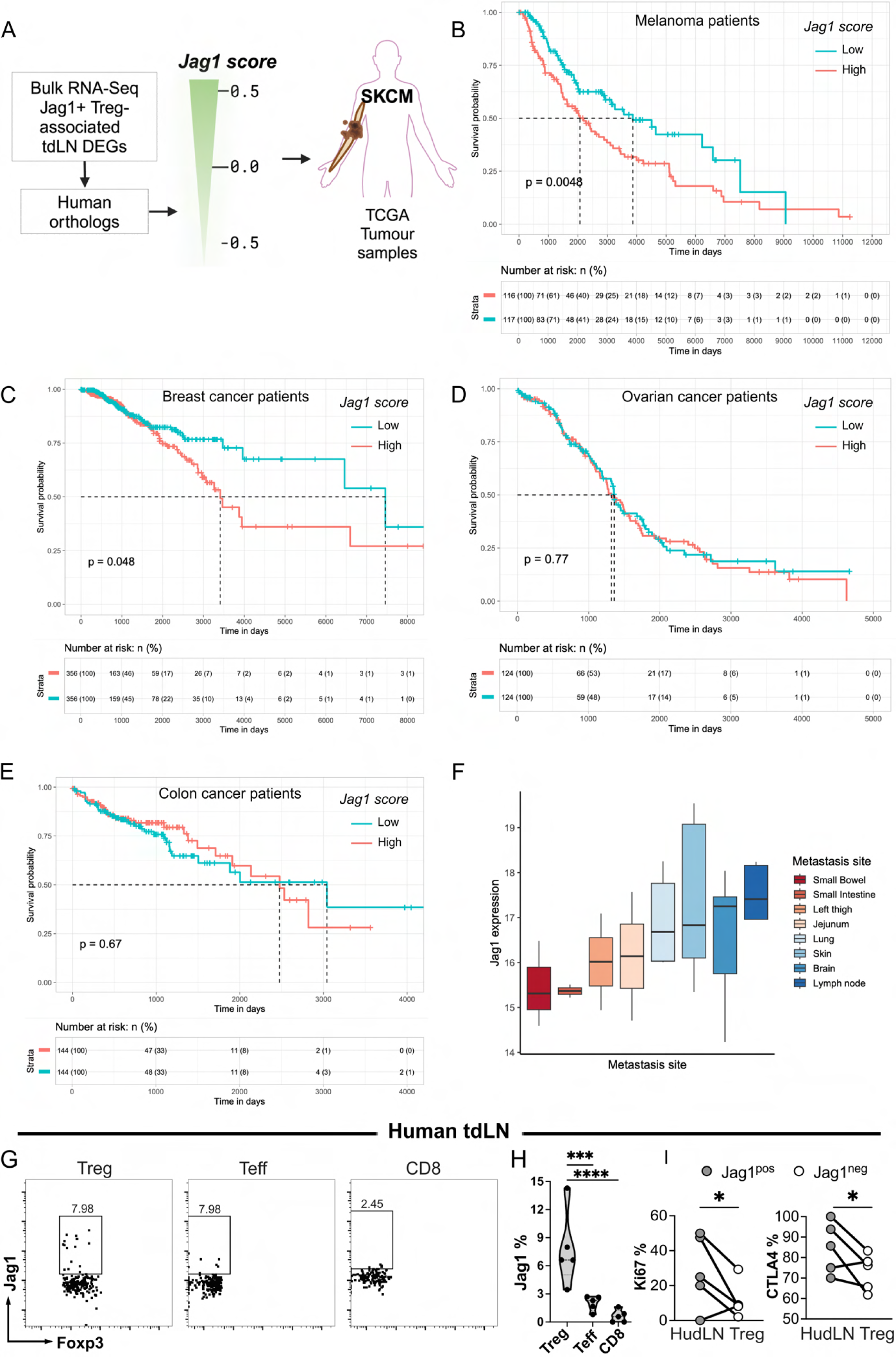
High Jag1 score on Tregs correlates with worse patient prognosis. **A)** Schematic outline of the TCGA exploitation workflow. Transcriptome signatures of Jag1^pos^ Tregs were extracted from the bulk tdLN RNA-Seq data and taken together to compute a Jag1-based scoring system. TCGA tumour samples were classified into low, medium and high, based on their Jag1 score, and tested for their association with patient overall survival. **B-E)** Graph illustrating the survival status of **B)** melanoma, **C)** breast, **D)** ovarian and **E)** colorectal cancer patients stratified by Jag1 score. **F)** Box plot illustrating the correlation between Jag1 expression and the likelihood of metastasis site for human melanoma. **G)** Flow cytometric plots to identify % expression of Jag1 on Tregs in HutdLNs. **H)** Quantification of percentage Jag1 expression on HutdLN Treg, Teff and CD8+ T cells. **I)** Flow cytometric quantification of percentage Ki67 and CTLA4 on Jag1^pos^ and Jag1^neg^ HutdLN Tregs. Statistics were calculated by Mantel Cox test for (B-E), by one-way ANOVA for (H), and by paired t-test for (I). ****p<0.0001, ***p<0.001, *p<0.05, ns p>0.05.

Given the propensity of Tregs to express Jag1 in Late stage tdLNs (**Figure 1C**), and their role in stromal cell-mediated remodelling of tdLNs (**Figure 5**), we next queried whether Jag1 expression strongly associated with metastasis site. Common sites of metastasis for melanoma include the skin, brain, lungs and liver.^33^ However, high overall Jag1 expression skewed metastasis potential towards lymph nodes (**Figure 6F**), likely partially promoted by augmented lymphangiogenesis.

To understand whether human Tregs also express Jag1, we characterised its expression in melanoma human patient tdLNs (HutdLNs) and healthy peripheral blood mononuclear cells (PBMCs) via flow cytometry (full gating strategy is shown in **Figure S7A**). HutdLNs specimens were obtained from five patients, and PBMC material was obtained from four healthy controls. **Table 1** provides an overview of the patient characteristics for individual HutdLNs. Flow cytometric analysis showed significantly enriched Jag1 expression in HutdLN Tregs relative to other T cell subtypes (**Figure 6G-H)**, as well as to healthy PBMC Tregs **(Figure S7B-C)**. Jag1^pos^ HutdLN Tregs but not Jag1^neg^ Tregs also expressed high levels of Ki67 and CTLA4 markers (**Figure 6I**), echoing findings from earlier murine Treg characterisation. Together, these data demonstrate that Jag1^pos^ Tregs are enriched in HutdLNs and represent a phenotypically activated and proliferative Treg subset that may serve as predictors of poor patient survival.

**Table 1.**
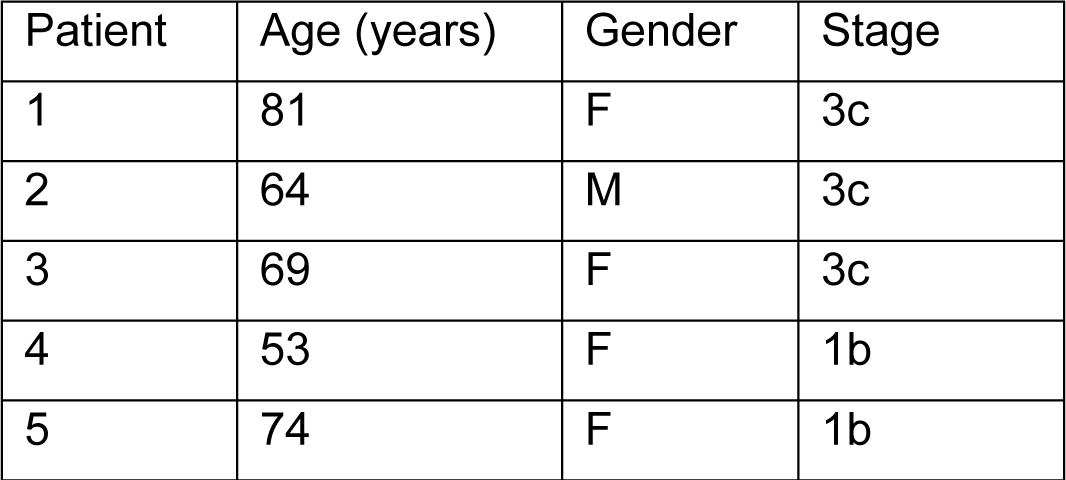
Patient characteristics of HutdLN specimens included in the study.

| Patient | Age (years) | Gender | Stage |
| --- | --- | --- | --- |
| 1 | 81 | F | 3c |
| 2 | 64 | M | 3c |
| 3 | 69 | F | 3c |
| 4 | 53 | F | 1b |
| 5 | 74 | F | 1b |

## DISCUSSION

Regulatory T cells (Tregs) traditionally play a central role in immune suppression, managing both innate and adaptive immune responses. Recent studies have highlighted additional non-immune functions of Tregs, including roles in tissue repair, angiogenesis, metabolism, and stem cell regeneration.^34^ A specialized subset of Tregs, characterized by Jag1 expression, has been identified as crucial for the regulation of the stem cell niche in the skin.^14^ Interestingly, Jag1 is globally overexpressed in several cancer types, including melanoma and breast cancer.^35,36^ In the latter, elevated protein levels can serve as an independent prognostic biomarker and thus a potential therapeutic target. High Jag1 expression is also significantly linked to lymph node metastasis and cancer recurrence.^37,38^ However, the role of Jag1 expressing Tregs in influencing the biological function of tissues other than skin, or during cancer progression, has not been investigated.

In the current study, we demonstrate that Jag1-expressing Tregs are present amongst the tumor-infiltrating Treg pool and exhibit an activated profile (Figure 1F-G). Notably, Jag1^pos^ Tregs were highly enriched in late tdLNs and exhibited an even higher degree of activation (Figure 1C-D). Key Treg-associated markers such as CD25, ICOS, and CTLA4 were upregulated in tdLN Jag1^pos^ Tregs, identifying Jag1 as a potential marker to delineate Treg activation status. Despite this phenotypic observation, the immunosuppressive capacity of Jag1^pos^ Tregs, whether in direct suppression of effector T cells or regulation of tissue inflammation, remains to be fully elucidated. Our immunophenotyping data showed no significant impact on the abundance and proliferation of tumour infiltrating and tdLN T cell subsets in this model (Figure 3), mirroring findings from skin studies where Jag1 was ablated in Tregs.^14^ These findings are highly suggestive that Jag1 expression in Tregs does not play a major role in influencing effector T cell activation *in vivo*.

Most strikingly, conditional deletion of Jag1 in Tregs restricted tdLN expansion and overall cellularity during late tumour progression. The largely uniform reduction in total cellularity across multiple immune and stromal cell subpopulations suggests a disruption in homeostatic cellular turnover in Foxp3^ΔJag1^ tdLNs. This could potentially be explained by altered lymphocyte homing or egress during LN priming. Immune profiling of endpoint tdLNs indicated no change in major T cell-mediated functional markers. Transcriptomic analysis of tdLNs indicated that Jag1^Pos^ Tregs control stromal-related markers, particularly those related to lymphatic endothelial cells (LECs). These cells line the sinuses of lymph nodes where they orchestrate formation of the highly organized lymphatic architecture. Remodelling of LN lymphatics occurs prior to metastatic colonization of tumour cells and involves the modulation of LECs in shaping the lymph node into pre-metastatic niches.^39^

Juxtacrine Jag1-Notch signalling between neighbouring endothelial cells is essential for vascular stability.^40^ Conditional Notch1 mutation in LECs result in pathological lymphangiogenesis and postnatal lymphatic defects.^41^ Similarly, endothelial lineage ablation of Jag1 negatively influences the central valve cellularity of the murine heart. Recent preliminary studies describe a role for Jag1 in promoting lymph node metastases in lymph node-invasive breast cancer via LEC mediated signalling.^38^ Our flow cytometric profiling of stromal subpopulations identified a significant loss of LECs in Foxp3^ΔJag1^ tdLNs compared to controls. Most notably, quantification of the two major hallmarks of lymphangiogenesis, lymphatic area and branching points of lymphatic vessels, were also were significantly dampened. Our data suggest a possible crosstalk between Tregs and LECs likely mediated by Notch signalling. It would be relevant to test if Jag1 loss in Tregs impacts LEC expression of endothelial markers regulating lymphocyte trafficking through lymph nodes, such as PLVAP and S1PR1.^42,43^ Evidence also suggests that Jag1 can potentially act as a secreted molecule in addition to direct cell-cell contact,^44^ making it difficult to decipher cell-cell interactions by colocalization analysis alone. Future studies, such as transwell co-culture assays, may uncover the requirement for any direct interactions.

Loss of Jag1 in Tregs disrupted lymphatic drainage in vivo, likely due to suppressed lymphangiogenesis, as evidenced by altered FITC-dextran fluorescence levels. Together, these observations support the notion that Jag1^pos^ Tregs may facilitate tdLN lymphangiogenesis and fluid drainage, processes known to promote metastasis to draining lymph nodes. Further investigation is needed to determine if Jag1^Pos^ Treg-induced lymphangiogenesis enhances metastatic potential. Additionally, the lymph antigenic load is largely specific to the type of tissue inflammatory response, and its transport to the draining lymph node is crucial for immunosurveillance. Whether the composition of lymph antigenic load is also impacted by Jag1 deletion remains to be determined.^45^

Exploitation of the TCGA database identified a positive correlation between high Jag1 score and poor overall survival in melanoma and breast cancer patients, but not in ovarian or colorectal cancer. A plausible explanation for this variability could be due to differences in immune cell infiltration, responsiveness to immune checkpoint inhibitors, or stromal content amongst cancer types.^46^ Skewed expression of Jag1 in Tregs compared to other T cell subsets in melanoma patient tdLNs suggests conservation of their role across species, highlighting the potential therapeutic benefits of targeting Jag1/Notch signaling in Tregs for melanoma progression.

Our Cre-flox model with systemic tamoxifen administration lacks tissue-specific modulation, complicating the elucidation of Treg functions in specific tissues. Recent advancements, such as skin-specific Treg manipulation techniques,^47^ could be adapted for other organs to achieve more precise modulation. Implementation of this topical application technique to other organs, such as a subcutaneously implanted B16 tumour, would enable more specific modulation of tumour infiltrating Tregs.

Additionally, our HutdLN stains showed sparse detection of Jag1^pos^ cells in the Treg gate, although still in greater proportions than other T cell subsets. This may be an accurate reflection of the true physiological circumstance or may be related to cell digestion and Treg recovery methods. Future experiments could include immunofluorescence or spatial transcriptomics to better understand the localization and abundance of Jag1^Pos^ Tregs, relative to Jag1^Neg^ Tregs or stromal cell populations, in patient tdLN tissue sections.

In conclusion, our study provides new insights into the function of Jag1^pos^ Tregs in tdLNs during tumor development. We propose that Jag1^pos^ Tregs promote lymphangiogenesis and establish a pro-tumorigenic environment within the primary tumor-draining lymph nodes, offering potential targets for therapeutic intervention.

## Acknowledgments

We thank the Advanced Cytometry Platform (Flow Core), Research and Development Department at Guy’s and St Thomas’ NHS Foundation Trust, and the Barts Cancer Institute Flow Cytometry Facility at Queen Mary University of London for assistance with flow cytometry experiments. Figures 1A, 2A, 6A and S6A were created using Biorender.com.

## Author contributions

Conceptualization, N.A.; methodology, N.A., J.Z.X., P.P.L., B.X., and A.E.D.; formal analysis, N.A., J.Z.X., and P.T.; investigation, J.Z.X., P.P.L., P.T., H.A., I.C., M.T.B., C.Z.L and H.Y.; data curation, J.Z.X., and K.S.; writing – original draft, N.A., and J.Z.X.; writing – review & editing, N.A., and J.Z.X.; visualization, N.A., J.Z.X., and P.T.; resources, A.E.D., and S.N.K.; supervision, N.A., and T.L.; funding acquisition, N.A.

## METHODS

### KEY RESOURCES TABLE

| REAGENT or RESOURCE | SOURCE | IDENTIFIER |
| --- | --- | --- |
| **Antibodies** |  |  |
| Human PBMC Panel |  |  |
| BUV395 Mouse Anti-Human CD3 | BioLegend | Cat# 563546, RRID: AB_2744387 |
| Brilliant Violet 785™ anti-human CD4 Antibody | BioLegend | Cat# 317442, RRID: AB_2563242 |
| Brilliant Violet 650™ anti-human CD8a Antibody | BioLegend | Cat# 301042, RRID: AB_2563505 |
| CD278 (ICOS) Antibody, anti-human/mouse/rat, VioGreen™, REAfinity™ | Miltenyi Biotec | Cat# 130-100-740, RRID: AB_2656919 |
| CD45 Antibody, anti-human, Vio® Bright B515, REAfinity™ | Miltenyi Biotec | Cat# 130-110-640, RRID: AB_2658255 |
| CD25 Antibody, anti-human, PE, REAfinity™ | Miltenyi Biotec | Cat# 130-113-286, RRID: AB_2733792 |
| CD127 Antibody, anti-human, PE-Vio® 770, REAfinity™ | Miltenyi Biotec | Cat# 130-113-415, RRID: AB_2733198 |
| Brilliant Violet 711™ anti-human Ki-67 Antibody | BioLegend | Cat# 350516, RRID: AB_2563861 |
| Alexa Fluor® 647 anti-human FOXP3 Antibody | BioLegend | Cat# 320214, RRID: AB_492984 |
| Brilliant Violet 605™ anti-human CD152 (CTLA-4) Antibody | BioLegend | Cat# 369610, RRID: AB_2632779 |
| FcR Blocking Reagent, human | Miltenyi Biotec | Cat# 130-059-901, RRID: AB_2892112 |
| Jagged1 Monoclonal antibody | Proteintech | Cat# 66890-1-Ig, RRID: AB_2882220 |
| Goat anti-Mouse IgG (H+L) Highly Cross-Adsorbed Secondary Antibody, Alexa Fluor™ Plus 594 | ThermoFisher | Cat# A32742, RRID: AB_2762825 |
| Mouse Stromal Cells Panel |  |  |
| CD45 Monoclonal Antibody (30-F11), Alexa Fluor™ 700, eBioscience™ | ThermoFisher | Cat# 56-0451-82, RRID: AB_891454 |
| Brilliant Violet 421™ anti-mouse Podoplanin Antibody | BioLegend | Cat# 127423, RRID: AB_2814017 |
| Brilliant Violet 711™ anti-mouse CD31 Antibody | BioLegend | Cat# 102449, RRID: AB_2860594 |
| FOXP3 Monoclonal Antibody (FJK-16s), FITC, eBioscience™ | ThermoFisher | Cat# 11-5773-80, RRID: AB_465243 |
| Ki-67 Monoclonal Antibody (SolA15), PE, eBioscience™ | ThermoFisher | Cat# 12-5698-80, RRID: AB_11149672 |
| Anti-Mouse CD16 / CD32 (Fc Sheild) | Tonbo Biosciences | Cat# 70-0161-M001 |
| TMA Staining Panels |  |  |
| Human/Mouse/Rat CD31/PECAM-1 Antibody | Biotechne/R&D System | Cat# AF3628, RRID: AB_2846942 |
| Anti-LYVE1 antibody - BSA and Azide free | Abcam | Cat# AB14917, RRID: AB_301509 |
| FOXP3 Monoclonal Antibody (FJK-16s), eBioscience™ | ThermoFisher | Cat# 14-5773-82, RRID: AB_467576 |
| CD31 (PECAM-1) (D8V9E) XP® Rabbit mAb | Cell Signaling Technology | Cat# 77699, RRID: AB_2722705 |
| Purified anti-mouse Podoplanin Antibody | BioLegend | Cat# 127401, RRID: AB_1089186 |
| Goat anti-Rabbit IgG (H+L) Highly Cross-Adsorbed Secondary Antibody, Alexa Fluor™ Plus 555 | ThermoFisher | Cat# A32732, RRID: AB_2633281 |
| Goat anti-Rat IgG (H+L) Highly Cross-Adsorbed Secondary Antibody, Alexa Fluor™ Plus 647 | ThermoFisher | Cat# A48265, RRID: AB_2895299 |
| Goat anti-Syrian Hamster IgG (H+L) Cross-Adsorbed Secondary Antibody, Alexa Fluor™ 488 | ThermoFisher | Cat# A-21110, RRID: AB_2535759 |
| Rhodamine Red™-X (RRX) AffiniPure™ Donkey Anti-Goat IgG (H+L) | Jackson ImmunoResearch | Cat# 705-295-147, RRID: AB_2340423 |
| Donkey anti-Rat IgG (H+L) Highly Cross-Adsorbed Secondary Antibody, Alexa Fluor™ 488 | ThermoFisher | Cat# A-21208, RRID: AB_2535794 |
| Donkey anti-Rabbit IgG (H+L) Highly Cross-Adsorbed Secondary Antibody, Alexa Fluor™ Plus 647 | ThermoFisher | Cat# A32795, RRID: AB_2762835 |
| Mouse T Cell Panel |  |  |
| CD45 Monoclonal Antibody (30-F11), Alexa Fluor 700, eBioscience™ | ThermoFisher | Cat# 56-0451-82, RRID: AB_891454 |
| CD3 Antibody, anti-mouse, PerCP-Vio® 700, REAfinity™ | Miltenyi Biotec | Cat# 130-120-826, RRID: AB_2752207 |
| TCRγ/δ Antibody, anti-mouse, PE-Vio® 770, REAfinity™ | Miltenyi Biotec | Cat# 130-123-290, RRID: AB_2802022 |
| Brilliant Violet 650™ anti-mouse CD4 Antibody | BioLegend | Cat# 100545, RRID: AB_11126142 |
| CD8a Antibody, anti-mouse, APC-Vio® 770, REAfinity™ | Miltenyi Biotec | Cat# 130-120-806, RRID: AB_2752203 |
| CD25 Antibody, anti-mouse, PE-Vio® 615, REAfinity™ | Miltenyi Biotec | Cat# 130-123-028, RRID: AB_2811434 |
| BD Horizon™ BV605 Hamster Anti-Mouse CD27 | BD Biosciences | Cat# 563365, RRID: AB_2738160 |
| Jagged1 Antibody (E-12) | Santa Cruz Biotechnology | Cat# Sc-390177, RRID: AB_2892141 |
| Brilliant Violet 711™ anti-human/mouse/rat CD278 (ICOS) Antibody | BioLegend | Cat# 313547, RRID: AB_2734288 |
| BD Horizon™ BV786 Mouse Anti-Ki-67 | BD Biosciences | Cat# 563756, RRID: AB_2732007 |
| FoxP3 Antibody, anti-mouse, Vio® R667, REAfinity™ | Miltenyi Biotec | Cat# 130-111-682, RRID: AB_2651771 |
| Brilliant Violet 421™ anti-mouse CD152 Antibody | BioLegend | Cat# 106311, RRID: AB_10901170 |
| Mouse Myeloid Panel |  |  |
| BD Horizon™ BUV395 Rat Anti-Mouse Ly-6G | BD Biosciences | Cat# 565964, RRID: AB_2716852 |
| Brilliant Violet 421™ anti-mouse I-A/I-E Antibody | BioLegend | Cat# 107632, RRID: AB_2650896 |
| Brilliant Violet 510™ anti-mouse/human CD45R/B220 Antibody | BioLegend | Cat# 103248, RRID: AB_2650679 |
| Brilliant Violet 510™ anti-mouse CD3 Antibody | BioLegend | Cat# 100234, RRID: AB_2562555 |
| Brilliant Violet 650™ anti-mouse CD11c Antibody | BioLegend | Cat# 117339, RRID: AB_2562414 |
| Brilliant Violet 785™ anti-mouse F4/80 Antibody | BioLegend | Cat# 123141, RRID: AB_2563667 |
| FITC anti-mouse Ly-6C Antibody | BioLegend | Cat# 128005, RRID: AB_1186134 |
| BD Pharmingen™ PerCP-Cy™5.5 Rat Anti-Mouse CD103 | BD Biosciences | Cat# 563637, RRID: AB_2738337 |
| CD207 (Langerin) Monoclonal Antibody (eBioL31), PE, eBioscience™ | ThermoFisher | Cat# 12-2075-82, RRID: AB_763452 |
| APC anti-mouse CD301b (MGL2) Antibody | BioLegend | Cat# 146813, RRID: AB_2566024 |
| CD45 Monoclonal Antibody (30-F11), Alexa Fluor 700, eBioscience™ | ThermoFisher | Cat# 56-0451-82, RRID: AB_891454 |
| APC/Cyanine7 anti-mouse/human CD11b Antibody | BioLegend | Cat# 101225, RRID: AB_830641 |
| **Experimental Models: Cell Lines** |  |  |
| B16F10 murine melanoma cells | Francis Crick Institute | N/A |
| **Experimental Models: Organisms/Strains** |  |  |
| WT C57BL/6J | Charles River | RRID: IMSR_CRL:027 |
| Foxp3<sup>eGFP-Cre-ERT2</sup> (Foxp3<sup>tm9(EGFP/cre/ERT2)Ayr/J</sup>) C57BL/6 | The Jackson Laboratory | RRID: IMSR_JAX:016961 |
| Jag1<sup>fl/fl</sup> (B6.129S-Jag1<sup>tm2Grid/J</sup>) | The Jackson Laboratory | RRID: IMSR_JAX:010618 |
| **Software and Algorithms** |  |  |
| FlowJo (version 10.8.1) | Becton Dickinson | https://www.flowjo.com/solutions/flowjo/; RRID:SCR_008520 |
| QuPath (version 0.5.0-arm64) | Bankhead et al.<sup>48</sup> | https://qupath.github.io/ |
| Prism (version 8.0) | GraphPad Software | http://www.graphpad.com/; RRID:SCR_002798 |
| Inkscape (version 1.3.2) | Inkscape | https://inkscape.org/; RRID:SCR_014479 |
| BioRender | BioRender | https://biorender.com/ |
| Trimmomatic (version 0.36) | Bolger, Lohse & Usadel, 2014<sup>49</sup> | http://www.usadellab.org/cms/index.php?page=Trimmomatic; RRID:SCR_011848 |
| STAR aligner (version 2.5.2b) | Dobin et al., 2013<sup>50</sup> | https://github.com/alexdobin/STAR |
| Subread (version 1.5.2) | Liao, Smyth & Shi, 2013<sup>51</sup> | https://subread.sourceforge.net/ |
| R (version 4.2.0) | CRAN | https://www.r-project.org/ |
| RStudio (version 2021.09.0) | RStudio | https://www.rstudio.com/ |
| R package: Seurat (version 5.0.1) | Gribov et al., 2010 | http://seurat.r-forge.r-project.org/; RRID:SCR_007322 |
| R package: DESeq2 (version 1.38.3) | Love et al., 2014 | https://bioconductor.org/packages/DESeq2; RRID:SCR_015687 |
| R package: EnhancedVolcano (version 1.16.0) | Blighe et al., 2023<sup>52</sup> | https://bioconductor.org/packages/EnhancedVolcano; RRID:SCR_018931 |
| R package: enrichR (version 3.2) | Chen et al., 2013<sup>53</sup> | https://maayanlab.cloud/Enrichr/; RRID:SCR_001575 |
| R package: pheatmap (version 1.0.12) | Kolde, 2019<sup>54</sup> | https://cran.r-project.org/web/packages/pheatmap/; RRID:SCR_016418 |
| R package: biomaRt (version 2.54.1) | Smedley et al., 2015<sup>55</sup> | http://www.biomaRt.org/; RRID:SCR_002987 |
| R package: singscore (version 1.18.0) | Foroutan et al., 2018<sup>56</sup> | https://github.com/DavisLaboratory/singscore/ |
| R package: survival (version 3.5.8) | Therneau, 2024<sup>57</sup> | https://cran.r-project.org/package=survival/; RRID:SCR_021137 |
| R package: survminer (version 0.4.9) | Kassambara, Kosinski & Biecek, 2021<sup>58</sup> | https://cran.r-project.org/web/packages/survminer/; RRID:SCR_021094 |
| R package: ggplot2 (version 3.5.0) | Wickham, 2016<sup>59</sup> | https://cran.r-project.org/web/packages/ggplot2/; RRID:SCR_014601 |
| **Other Datasets** |  |  |

## CONTACT FOR REAGENT AND RESOURCE SHARING

Requests for reagents and resources should be directed to the Lead Contact, Niwa Ali

## EXPERIMENTAL MODEL AND SUBJECT DETAILS

### Cell lines

The B16F10 melanoma cell line was obtained from Cell Services STP at The Francis Crick Institute and cultured in RPMI 1640 (Thermo Fisher; 21875-034) supplemented with 10% FBS (ThermoFisher; 10270-106) and 1% Pen-Strep (Sigma-Aldrich; P4458-100ML). Cells were maintained in a 5% CO_2_ incubator at 37°C and passaged every 2-3 days.

### Mice

Wild-type, Foxp3^Cre-ERT2^Jag1fl/fl and fl/wt mice of B6 background were fed with a standard chow diet and housed in line with UK regulations. Husbandry of animals were performed by trained animal facility staff in the New Hunt’s House Biological Services Unit at KCL. All experiments were performed on animals with no prior procedures, with authorisation under the project license PP6051479. Littermates of the same sex were genotyped and assigned to experimental groups based on genotyping results. For conditional Jag1 knockout in the syngeneic tumour model, 6-8 week old Foxp3^eGFP-Cre-ERT2^Jag1fl/fl and fl/wt female mice received 5 consecutive doses of tamoxifen (Sigma-Aldrich; T5648) dissolved in corn oil at 0.1mg per gram of body weight. For conditional Jag1 knockout in the skin wounding model, 6-8 week old Foxp3^eGFP-Cre-ERT2^Jag1fl/fl and fl/wt female mice received 4 doses of tamoxifen at 0.1mg per gram of body weight prior to dorsal back wounding. Two more doses were given upon wounding prior to harvest.

Syngeneic tumour model 2 × 10^5^ B16F10 tumour cells mixed with matrigel at 1:1 ratio were intradermally injected into the left flank of 6-11 week old Foxp3-DTR (Jackson Laboratory) and Foxp3^eGFP-Cre-ERT2^Jag1fl/fl or fl/wt female mice. Tumour size was measured by digital caliper every 3 days and animals were sacrificed and lymph nodes and tumours were collected at various time points. Endpoint tissues were weighed and tumour volumes was calculated using the formula (π/6)(length*width*height). Alternatively, tumour volumes prior to endpoint were predicted using (1/2)(length*width^2^).

### FITC-dextran permeability assay *in vivo*

Endpoint tumour-bearing mice received either 5mg/ml 4 kDa (Sigma-Aldrich; 46944) or 70 kDa (Sigma-Aldrich; 46945) FITC-dextran in PBS subcutaneously in the left footpad 10 minutes prior to culling. Lymph nodes and tumours were collected and homogenized with CK14 soft tissue lysing beads (Precellys/Bertin; P000912-LYSK0-A) in 1 ml PBS. Supernatant was transferred to flat-bottom 96 well plate and the blue 475 nm excitation profile was ran on the GloMax® Discover fluorescence plate reader (Promega) to detect FITC emission at 500-550 nm.

### Skin wounding assay

Mice were anaesthetized, shaved and injected with Vetergesic. Full-thickness wounds were created on the dorsal back with 4mm biopsy punches (Stifel Laboratory Research). Wound-dLNs were collected at 5dpw for flow cytometry (and processed as described below).

## METHOD DETAILS

### Tissue digestion

For T cell immunophenotyping, lymph nodes were passed through 70 µm cell strainers and B16F10 tumour tissues were minced then digested with 1 ml mixture of 2 mg/ml Collagenase (Sigma-Aldrich; C9407-1G), 0.1 mg/ml DNase I (Sigma-Aldrich; DN25-500MG) in 10% FBS RPMI1640 media per 200 mg tumour at 37°C for 45 minutes with 255 rpm rotation. Digested tumours were washed then passed through 100 µm followed by 40 µm strainers to retrieve single cell suspension. For stromal cell phenotyping, lymph nodes were digested using a protocol adapted from Fletcher et al., 2011. Briefly, lymph nodes were pierced with a 25g needle, then immersed in 0.5 ml digestion buffer of 0.2 mg/ml Collagenase P (Roche; 11213857001), 0.1 mg/ml DNase I (Roche; 10104159001), 0.8 mg/ml Dispase II (Roche; 04942078001) per lymph node at 37°C for 15 minutes. Eppendorf tubes each containing one lymph node were inverted every 5 minutes and after the final inversion, lymph nodes with triturated using 1ml pipette tips before supernatant containing released cells was collected with FACS-EDTA (2% FBS, 2mM EDTA (Thermo Fisher; 15575020) in PBS) buffer. The digestion process was repeated twice more by adding fresh 0.5 ml digestion buffer, incubating and collecting supernatant to the collection tubes. Finally, digested cells were spun down at 400g for 5 minutes and filtered through 70 µm strainers to remove collagen and debris. All cells were counted using a NC-200 NucleoCounter (Chemometec) and plated on round-bottom plates for staining.

Compensation bead controls were prepared by adding one drop of ArC™ reactive bead (Invitrogen; A10346) or UltraComp eBeads™ (Invitrogen; 01-2222-42) or both MACS® Comp bead - blank and anti-REA (Miltenyi Biotec; 130-104-693) to each well. Beads were stained with the corresponding antibody at the same dilution for 20 minutes on ice in the dark and fixed following the same protocol for tissue samples.

### Mouse tissue flow cytometry

For T cell immunophenotyping, single cell suspensions were stained with Zombie UV^TM^ Live/Dead (Biolegend; 423107) in PBS for 20 minutes at 4°C then washed with FACS buffer (2% FBS 2mM EDTA in PBS) and centrifuged at 1800 rpm for 4 minutes. Cells were then stained with surface antibodies resuspended in brilliant buffer (BD Biosciences; 566348) for 20 minutes at 4°C. Cells were washed, fixed with fixation buffer provided in the Foxp3 staining buffer kit (eBioscience; 00-5523-00) and stained with intracellular antibodies resuspended in the provided permeabilisation buffer for 20 minutes at 4°C. For stromal cell phenotyping, all steps were same as above except that cells were stained with surface antibodies for 60 minutes, fixed with fixation buffer for 30 minutes and stained with intracellular antibodies overnight, all at 4°C. Fluorescent conjugated antibodies used are listed in Key Resources Table. Flow cytometry was performed on the LSRFortessa II (BD Biosciences) analyzer. Standardization was performed between experiments using SPHERO^TM^ Rainbow calibration particles (BD Biosciences; 559123). Relevant gatings and data analysis was performed with the FlowJo software v.10.8.1.

### Tissue microarray (TMA) and immunofluorescence

Lymph node tissues were fixed in 10% neutral buffered formalin (NBF; Fisher Scientific; 10270219) overnight at 4°C. On the next day, fixed lymph nodes were washed with PBS twice on a rocking platform. Upon washes, fixed tissues were transferred into 70% Ethanol overnight before dehydration and embedding in paraffin. Individual paraffin-embedded tissue cores (1 mm diameter) were made into a recipient TMA block with a tissue arrayer (Beechers Instruments). The recipient block was populated with murine endpoint B16F10 tdLN samples (*N* = 28; *n* = 14; *N* = number of mice, *n* = number of cores analysed). Tissue arrays were sectioned at 4 µm for immunostaining. Sections were baked at 60°C for 30 minutes, deparaffinized with xylene then rehydrated with ethanol of descending concentrations. Following rehydration, sections were permeabilized with 0.05% Triton-X in TBS-T for 15 minutes at room temperatures. Sections were washed then heated in 0.01 M sodium citrate buffer (pH 6.0) at 96°C for 30 minutes for antigen retrieval. Unspecific epitopes were blocked with 30 minutes incubation in blocking solution (0.25 M glycine in 5% FBS TBS solution) at room temperature. Subsequently, sections were stained with primary antibodies diluted in 5% FBS TBS solution overnight at 4°C. Washed sections were then incubated in secondary antibodies diluted in 5% FBS TBS solution for 1 hour at room temperature. Finally, 300 nM DAPI solution was added to sections for 10 minutes at room temperature. Microscopy was performed using the VS120 fluorescent slide scanner (Olympus) with a × 20 objective. Fluorescent images were analyzed using QuPath v.0.5.0. Gaussian thresholding was applied to filter for CD31+ PDPN-areas corresponding to blood vessels, and CD31+ PDPN+ (or LYVE1+) areas corresponding to lymphatic vessels. Area of blood and lymphatic vessels was measured per ROI. PDPN MFI intensity was quantified by measuring PDPN single channel intensity in CD31+ PDPN+ areas per ROI. Length of lymphatic vessels was determined by measuring the distance from one end to the other end of lymphatic vessels per ROI.

### Mouse tissue RNA extraction

Lymph node and tumour tissues (≤ 5mg) were protected in RNAlater (Sigma; R0901-500ML) at −20°C until being homogenized with CK14 lysing beads in buffer RLT provided in RNeasy® Micro kit (Qiagen; 74034). Total RNA was extracted following the manufacturer’s protocol. RNA quality was checked using the TapeStation system (Agilent), only RNA samples with RIN score over 9.0 were sent to Genewiz for Illumina NovaSeq 2×150 bp paired-end mRNA sequencing.

### Human tumour and sentinel LN tissue flow cytometry

Liquid nitrogen frozen human sample vials were thawed and rested in 10ml of 10% FBS RPMI 1640 media containing 1MU/ml of DNase I (Sigma-Aldrich; 260913-25MU) for 1 hour in 37°C 5% CO_2_ incubator. Upon resting, 2 million cells per sample were added with FcR blocking reagent (Miltenyi Biotec; 130-059-901) diluted in FACS buffer (5% FBS in PBS) for 20 minutes at 4°C then washed with FACS and centrifuged at 300 g for 5 minutes. Cells were then stained with Zombie UV^TM^ Live/Dead for 20 minutes at 4°C. Next, cells were stained with a purified primary Jag1 antibody (Proteintech) resuspended in FACS buffer for 30 minutes at room temperature, then counter-stained with a goat anti-mouse AF594 secondary antibody (Thermo Fisher) resuspended in FACS buffer for 15 minutes at room temperature. Washed cells were then stained with a mixture of surface antibodies resuspended in brilliance buffer for 20 minutes at 4°C. Cells were then fixed with 1 ml of fixation buffer provided in the Foxp3 staining buffer kit for 30 minutes. During fixation, cells were vortexed for 10 seconds every 10 minutes. Fixed cells were washed twice with 1 ml permeabilization buffer and centrifuged at 800 g for 5 minutes before being stained with intracellular antibodies resuspended in the permeabilisation buffer for 2 hours at 4°C. Finally, cells were washed once with 1 ml of permeabilization buffer and once with 1 ml of PBS before being ready for flow cytometry. Purified, secondary and fluorescent conjugated antibodies used here are listed in Key Resources Table.

## QUANTIFICATION AND STATISTICAL ANALYSIS

Parameters such as sample size, dispersion or precision are reported in Figure Legends. Statistical analyses were performed in Prism 8.0 (GraphPad). Details of the statistics and appropriate test used are also indicated in Figure Legends. *p < 0.05, **p < 0.01 ***p < 0.001, ****p < 0.001, p-values greater than 0.05 was identified as not statistically significant.

### Whole tissue bulk RNA-Sequencing (RNA-Seq) analysis

Sequence reads were trimmed using Trimmomatic v.0.36 and mapped to the Mus musculus GRCm38 genome by STAR aligner v2.5.2b. Unique gene hits that fell within exon regions were counted from generated BAM files using the Subread package v.1.5.2. Differential expression analysis was performed on pre-processed count table with the “DESeq2” package v.1.38.3. Volcano plots highlighting a subset of significant differentially expressed genes (DEGs) with absolute log_2_ fold change >1 and adjusted p-value < 0.05 were plotted by the “EnhancedVolcano” package v.1.16.0. Gene ontology (GO) analysis was performed to extract enriched biological pathways in the Reactome 2022 database using the “enrichR” package v.3.2. A heatmap of A specific pathway of interest was created using the “pheatmap” package v.1.0.12. Upregulated and downregulated DEGs of Jag1^pos^ vs. Jag1^neg^ tumour-draining lymph nodes (tdLN) were converted to human gene orthologs with “biomaRt” v.2.54.1. The “singscore” method v.1.18.0 was used to score individual samples by their Jag1^pos^ score based on the Jag1 differential gene set provided.

### TCGA data exploitation

Count data was normalized using “DESeq2” and a Jag1^pos^ score was calculated per sample using the simpleScore function in singscore. Subsquently, tumour patients were classified into 3 groups (Low, Medium and High), based on the Jag1^pos^ score. The “survival” package v.3.5.8 was used to perform Cox regression analysis on survival data. Later, the “survminer” package v.0.4.9 was used to construct Kaplan-Meier curves.

### Public mouse single cell RNA-sequencing (scRNA-Seq) data analysis

The publicly available scRNA-Seq data of day 5, 8 and 11 CD45^+^ and CD45^-^ cells from B16F10 melanoma tumour and tdLN tissues was downloaded from https://www.ebi.ac.uk/gxa/sc/experiments/E-EHCA-2/Results. The “Seurat” package v.5.0.1 was used to log(x+1)-transformed and scale count data. tdLN data was extracted and plotted as a UMAP to show temporal changes in proportion of immune and non-immune populations based on author-inferred cell types. Using the “ggplot2” package v.3.5.0, a frequency stacked bar chart was also plotted to show frequency changes of cell populations quantitatively. In addition, WT-Upregulated DEGs from lymph node bulk RNA-sequencing were mapped onto this scRNA-Seq dataset to infer cell-type specificity.

## DATA AND CODE AVAILABILITY

Bulk RNA-Seq data of tumours and tdLNs will be deposited in ArrayExpress. Bulk RNA-Seq data of sorted Jag1^pos^ and Jag1^neg^ Tregs from tdLNs has been deposited in ArrayExpress. Other data is available from the corresponding author on reasonable request. Gene expression and survival data of melanoma (SKCM), breast cancer (BRCA), ovarian cancer (OV) and colon cancer (COAD) were downloaded UCSC Xena Browser. The publicly available scRNA-Seq data of day 5, 8 and 11 CD45^+^ and CD45^-^ cells from B16F10 melanoma tumour and tdLN tissues was downloaded from https://www.ebi.ac.uk/gxa/sc/experiments/E-EHCA-2/Results.

## Supplementary Figures

**Figure S1.**
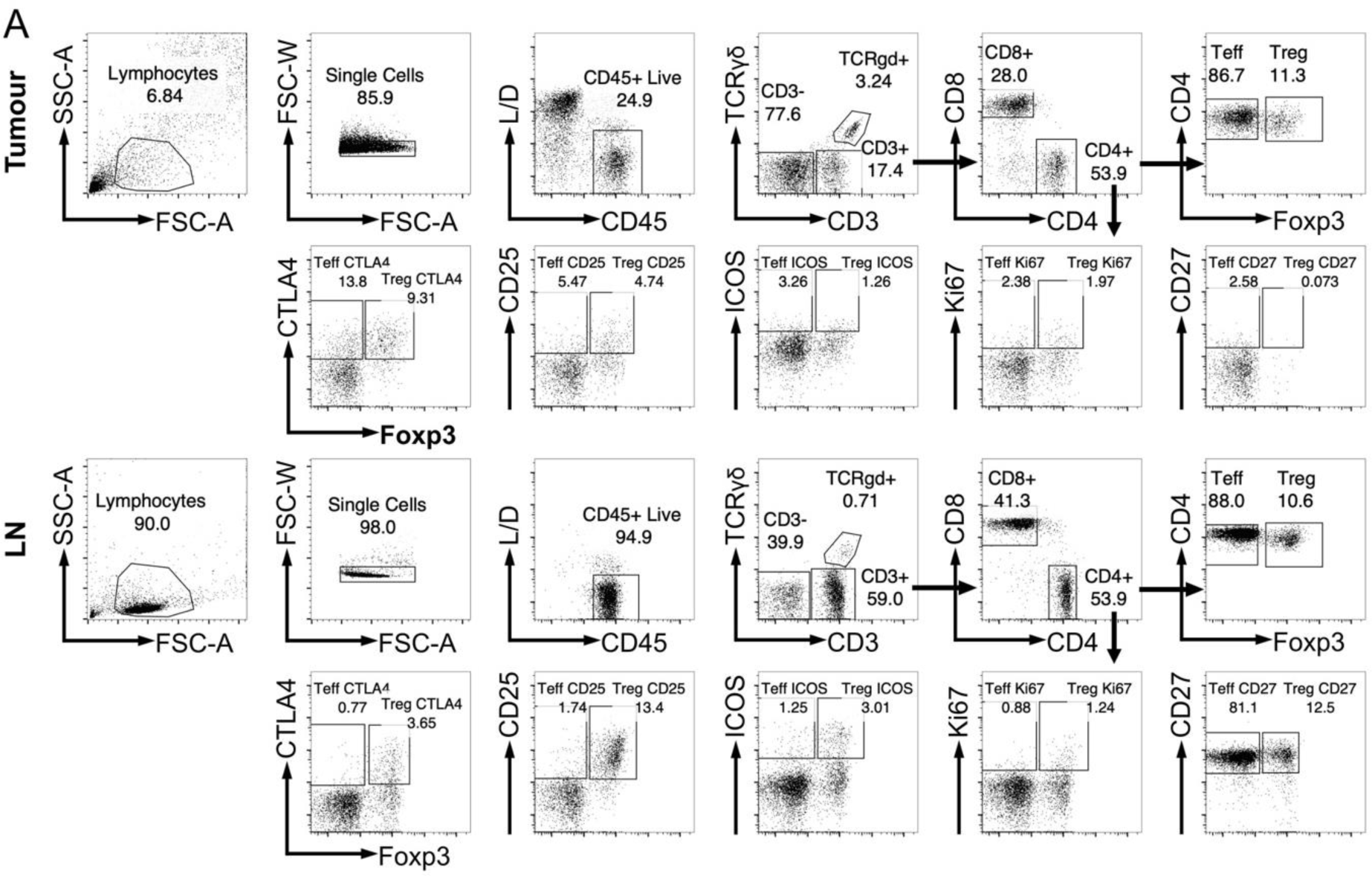
Flow cytometric gating strategy to identify % expression of Jag1, CTLA4, CD25, ICOS, Ki67 and CD27 on Tregs in tumors and LNs. All gatings were determined by their corresponding FMOs.

**Figure S2.**
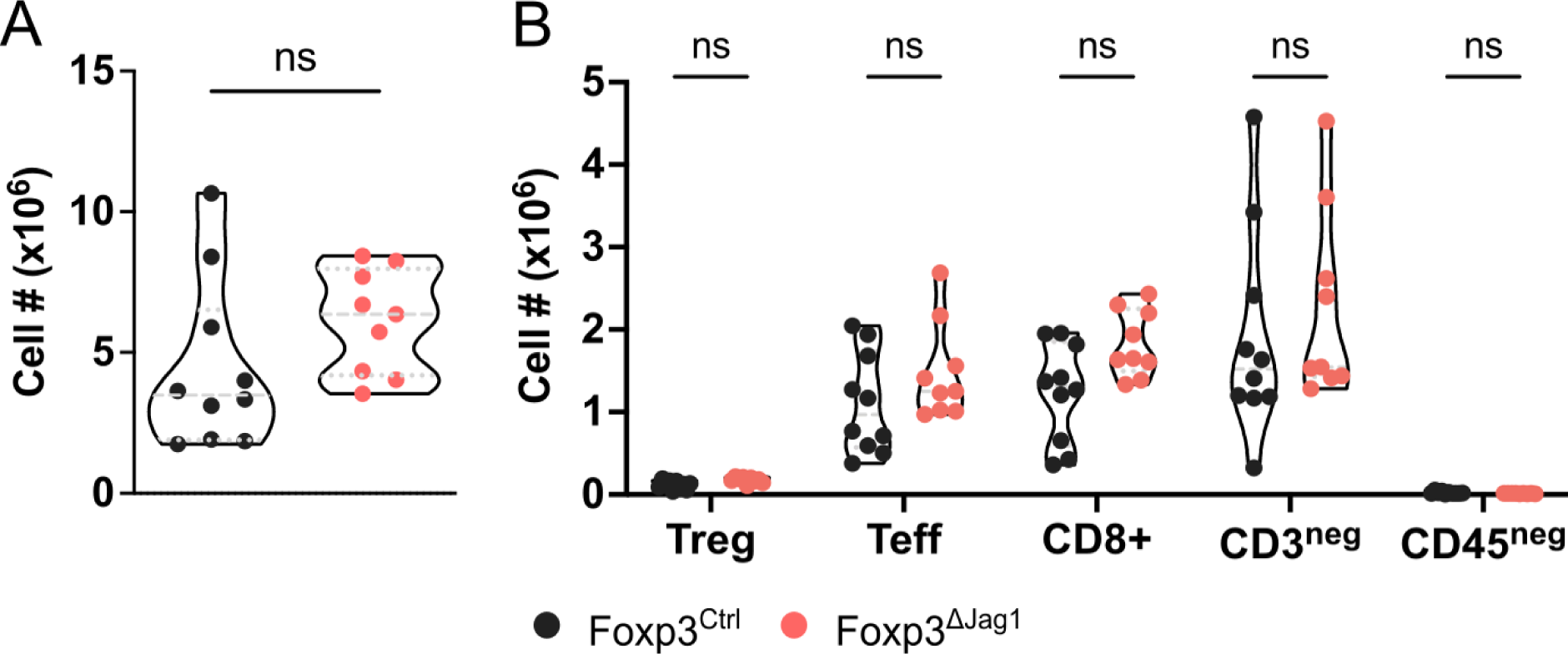
Loss of Jag1 in Tregs has no impact on wound-draining lymph node expansion. Foxp3^ΔJag1^ and Foxp3^Ctrl^ mice received full thickness wounds on the dorsal skin. **A)** Absolute number quantification of total cells in wound draining lymph nodes and **B)** Treg, Teff, CD8+ T cell, CD3- and CD45-subpopulations between Foxp3^ΔJag1^ and Foxp3^Ctrl^ tdLNs. Data are representative of two independent experiments. Statistics were calculated by unpaired t-test in (A), and by multiple unpaired t-test for (B). ns p>0.05.

**Figure S3.**
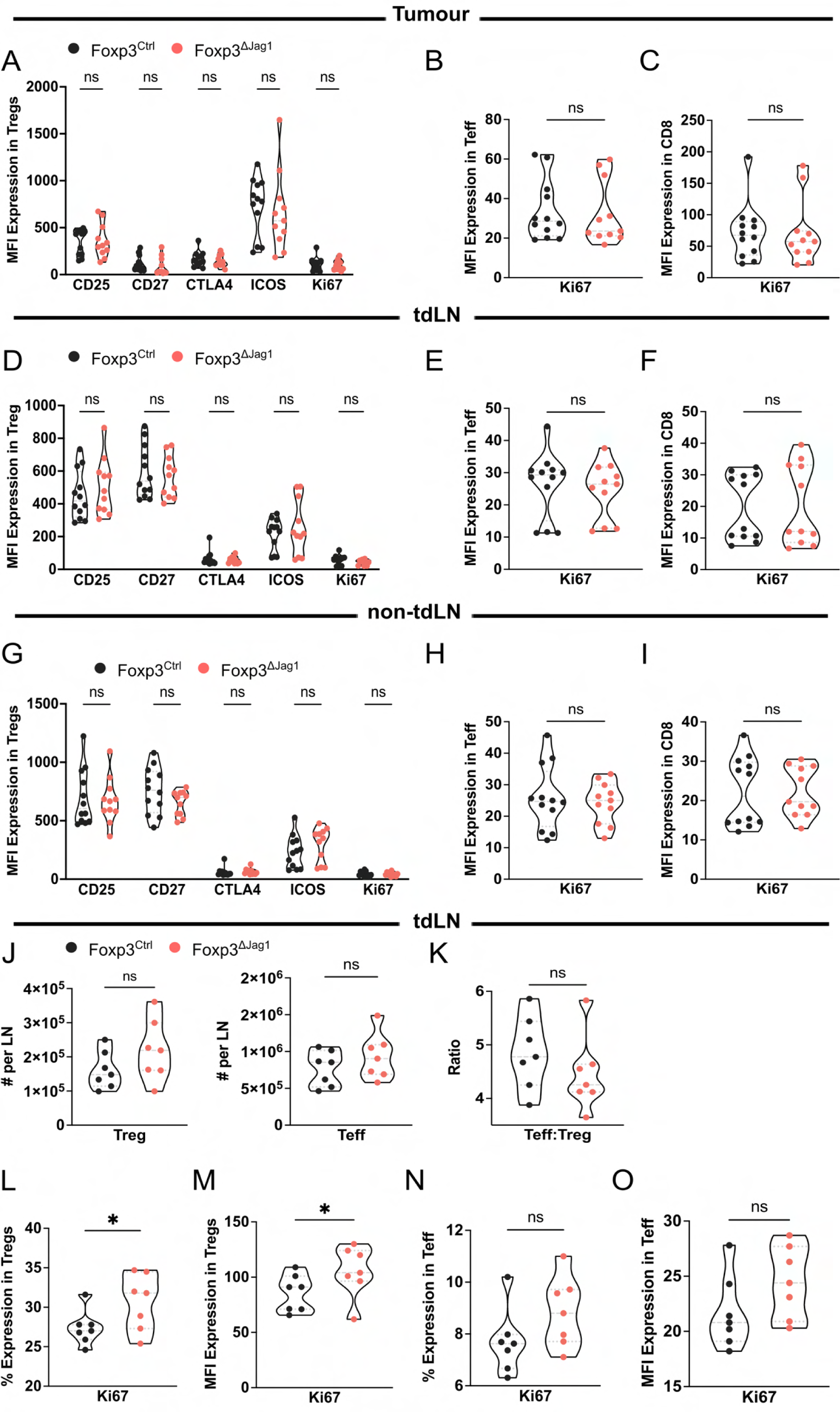
Jag1^pos^ Tregs do not influence immediate T cell-mediated anti-tumour immunity. **A)** Flow cytometric quantification of MFI values of activation markers CD25, CD27, CTLA4, ICOS and KI67 on endpoint Foxp3^ΔJag1^ and Foxp3^Ctrl^ tumour Tregs. **B-C)** MFI Ki67 expression on endpoint Foxp3^ΔJag1^ and Foxp3^Ctrl^ tumour **B)** Teffs and **C)** CD8+ T cells. **D)** Flow cytometric quantification of MFI values of activation markers on endpoint Foxp3^ΔJag1^ and Foxp3^Ctrl^ tdLN Tregs. **E-F)** MFI Ki67 expression on endpoint Foxp3^ΔJag1^ and Foxp3^Ctrl^ tdLN **E)** Teffs and **F)** CD8+ T cells. **G)** Flow cytometric quantification of MFI values of activation markers on endpoint Foxp3^ΔJag1^ and Foxp3^Ctrl^ non-tdLN Tregs. **H-I)** MFI Ki67 expression on endpoint Foxp3^ΔJag1^ and Foxp3^Ctrl^ non-tdLN **H)** Teffs and **I)** CD8+ T cells. **J)** Absolute number quantification of Tregs and Teffs in D4 Foxp3^ΔJag1^ and Foxp3^Ctrl^ tdLNs. **K)** Teff:Treg and CD8:Treg ratios between D4 Foxp3^ΔJag1^ and Foxp3^Ctrl^ tdLNs. **L-M)** Flow cytometric quantification of **L)** percentage and **M)** MFI Ki67 expression on D4 tdLN Tregs. **N-O)** Flow cytometric quantification of **N)** percentage and **O)** MFI Ki67 expression on D4 tdLN Teffs. Data are representative of three independent experiments. Statistics were calculated by unpaired t-test in (A, D, G), and by multiple unpaired t-test for (B-C, E-F, H-O). *p<0.05, ns p>0.05.

**Figure S4.**
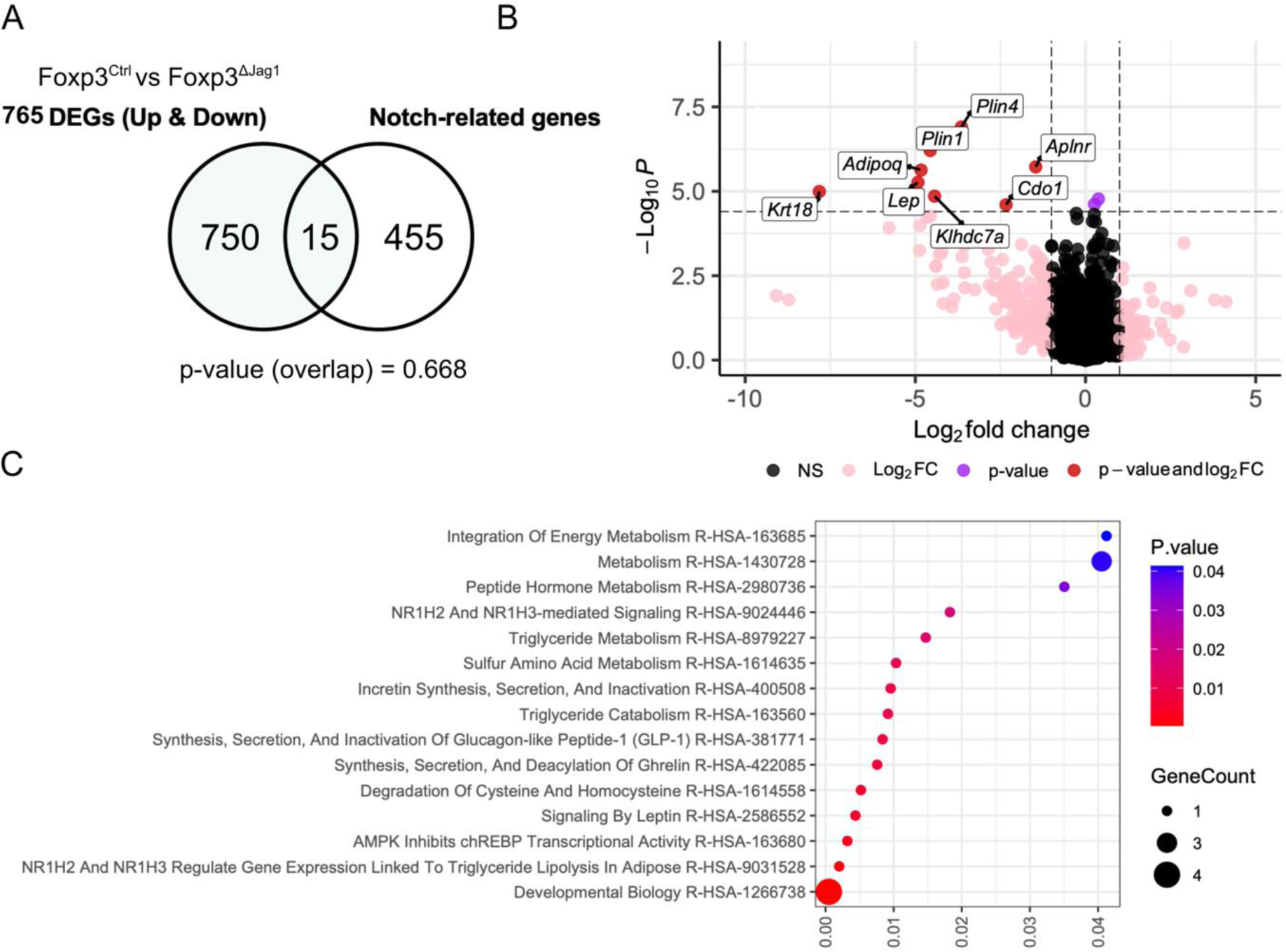
Loss of Jag1 on Tregs minimally impacts the tumour transcriptome. **A)** Venn diagram showing the overlap between tumour bulk RNA-Seq total DEGs (p-value <0.05) and known Notch-related genes. P-value represents the significance of the overlap as calculated by chi-squared test. **B)** Differential gene analysis between Foxp3^ΔJag1^ and Foxp3^Ctrl^ tumours. A negative fold change denotes Foxp3^Ctrl^-upregulated genes and a positive fold change denotes Foxp3^Ctrl^-downregulated genes. Dots in red are individual DEGs with >2 log2 fold change and <0.05 p-adjusted value. **C)** Reactome pathway enrichment analysis using tumour Foxp3^Ctrl^-upregulated DEGs (p-adjusted value <0.05).

**Figure S5.**
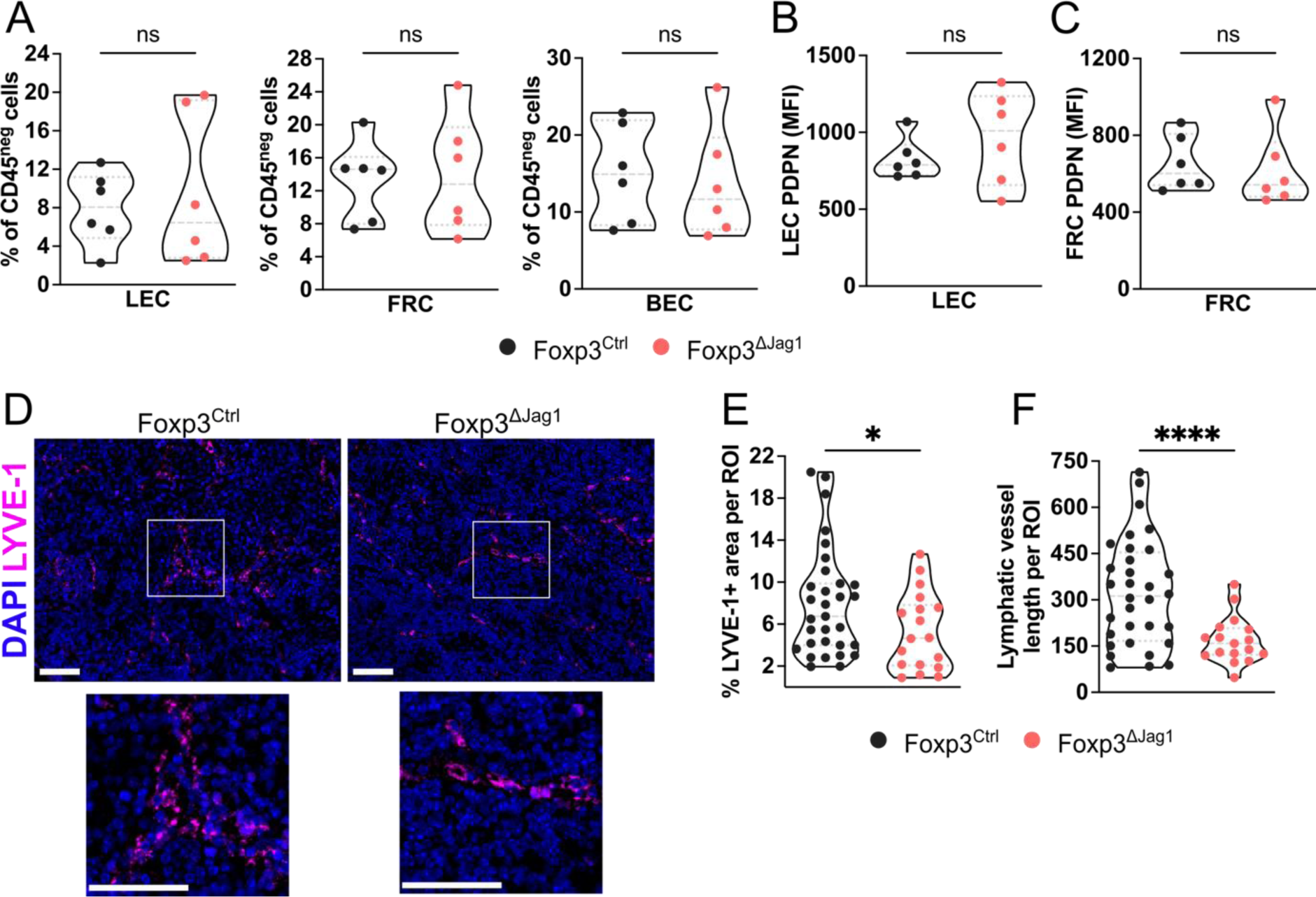
The proportion and PDPN MFI of LECs are unaffected in Day 4 Foxp3^ΔJag1^ tdLNs. **A)** Flow cytometric quantification of the percentage of stromal subpopulations (LECs, FRCs, BECs) out of total CD45-cells between Day 4 Foxp3^ΔJag1^ and Foxp3^Ctrl^ tdLN. **B-C)** Quantification of MFI PDPN on **B)** LECs and **C)** FRCs in Day 4 Foxp3^ΔJag1^ and Foxp3^Ctrl^ tdLN. **D)** Immunofluorescence staining of LYVE-1+ lymphatic vessels in Foxp3^ΔJag1^ and Foxp3^Ctrl^ endpoint tdLNs. 50 µm scale bar. **E)** % LYVE-1+ area and **F)** lymphatic vessel length quantification per ROI of Foxp3^ΔJag1^ and Foxp3^Ctrl^ tdLNs at endpoint. Data are representative of three independent experiments. Statistics were calculated by unpaired t-test. ***p<0.001, *p<0.05, ns p>0.05.

**Figure S6.**
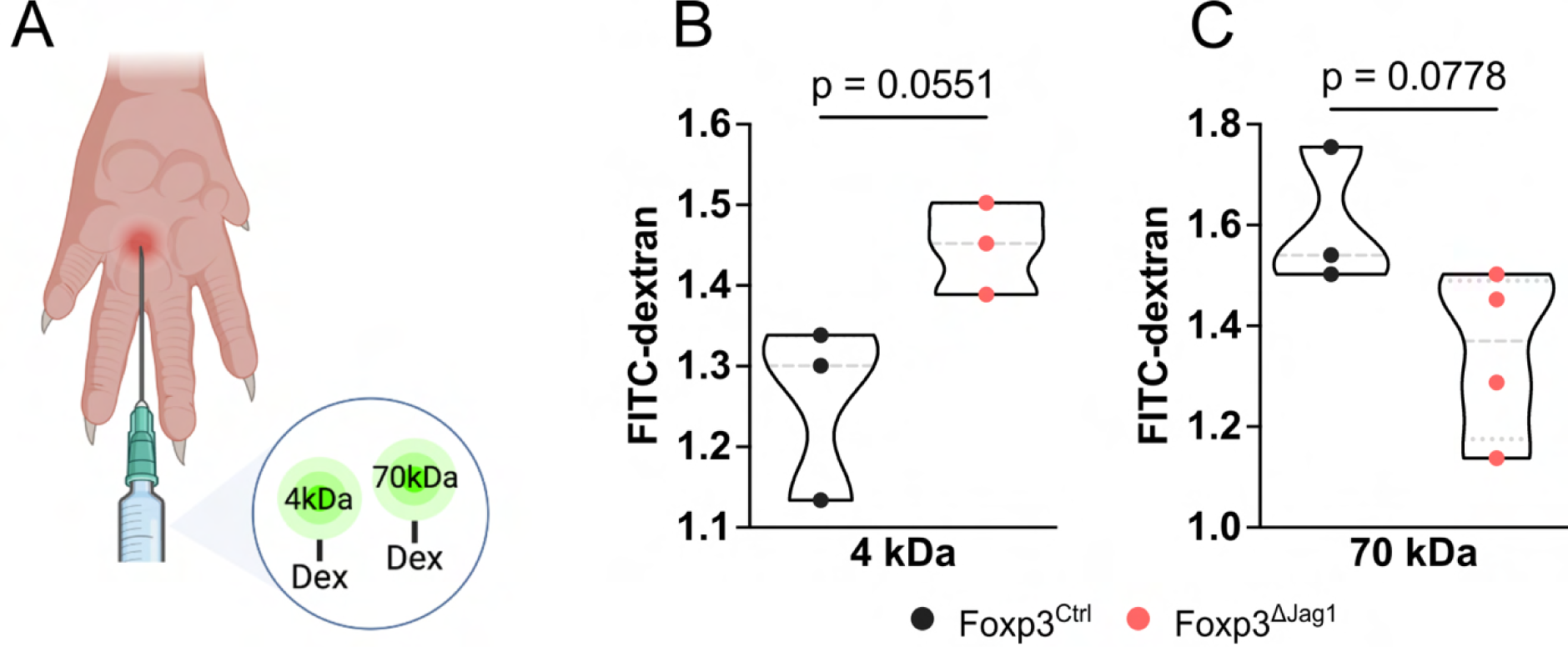
Jag1^pos^ Tregs modulate lymphatic drainage to tdLNs. **A)** Schematic outline of *in vivo* FITC-dextran assay to study LN drainage. **B-C)** Quantification of the fold change in **B)** 4kDa dextran FI and **C)** 70kDa dextran FI relative to FITC-negative LN control. Data are representative of 2 independent experiments. Statistics were calculated by unpaired t-test. Calculated p values are shown.

**Figure S7.**
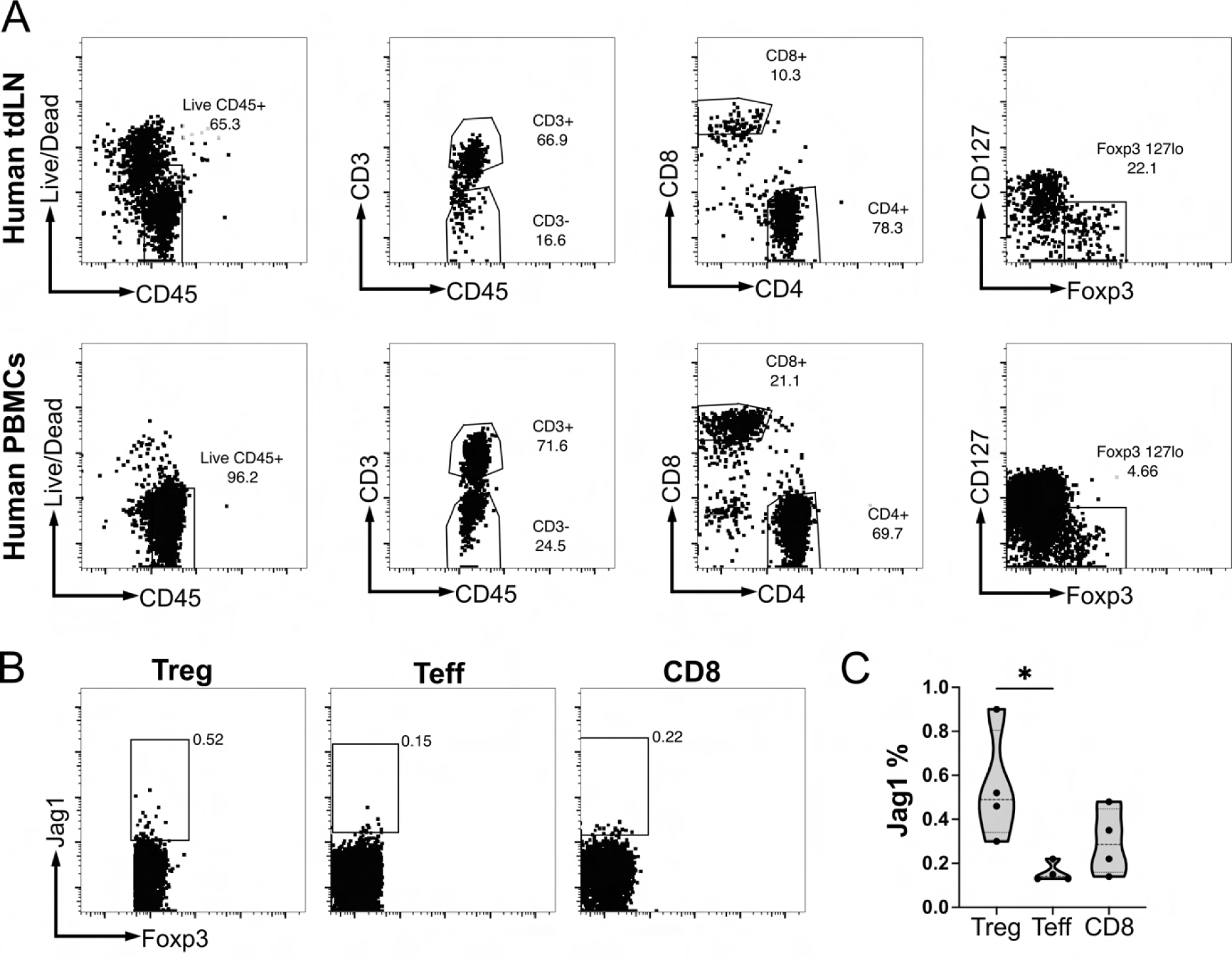
Jag1 is minimally expressed on PBMC T cell subpopulations. **A)** Flow cytometric gating strategy to identify Tregs, Teffs and CD8+ T cells in HutdLNs and PBMCs. All gatings were determined by their corresponding FMOs. **B)** Flow cytometric plots to identify % expression of Jag1 on Tregs in PBMCs. **C)** Quantification of percentage Jag1 expression on PBMC Tregs, Teffs and CD8+ T cells. Statistics were calculated by one-way ANOVA for (C). *p<0.05, ns p>0.05.

## BIBLIOGRAPHY

1. Ohue, Y., and Nishikawa, H. (2019). Regulatory T (Treg) cells in cancer: Can Treg cells be a new therapeutic target? Cancer Sci 110, 2080. 10.1111/CAS.14069.

2. du Bois, H., Heim, T.A., and Lund, A.W. (2021). Tumor-Draining Lymph Nodes: At the Crossroads of Metastasis and Immunity. Sci Immunol 6, eabg3551. 10.1126/SCIIMMUNOL.ABG3551.

3. Murthy, V., Katzman, D.P., Tsay, J.C.J., Bessich, J.L., Michaud, G.C., Rafeq, S., Minehart, J., Mangalick, K., de Lafaille, M.A.C., Goparaju, C., et al. (2019). Tumor-draining lymph nodes demonstrate a suppressive immunophenotype in patients with non-small cell lung cancer assessed by endobronchial ultrasound-guided transbronchial needle aspiration: A pilot study. Lung Cancer 137, 94–99. 10.1016/J.LUNGCAN.2019.08.008.

4. Frijlink, E., Bosma, D.M.T., Busselaar, J., Battaglia, T.W., Staal, M.D., Verbrugge, I., and Borst, J. (2024). PD-1 or CTLA-4 blockade promotes CD86-driven Treg responses upon radiotherapy of lymphocyte-depleted cancer in mice. Journal of Clinical Investigation 134. 10.1172/JCI171154.

5. Wang, Y., Zhu, T., Shi, Q., Zhu, G., Zhu, S., and Hou, F. (2024). Tumor-draining lymph nodes: opportunities, challenges, and future directions in colorectal cancer immunotherapy. J Immunother Cancer 12, e008026. 10.1136/JITC-2023-008026.

6. Commerford, C.D., Dieterich, L.C., He, Y., Hell, T., Montoya-Zegarra, J.A., Noerrelykke, S.F., Russo, E., Röcken, M., and Detmar, M. (2018). Mechanisms of Tumor-Induced Lymphovascular Niche Formation in Draining Lymph Nodes. Cell Rep 25, 3554–3563.e4. 10.1016/j.celrep.2018.12.002.

7. Jang, J.Y., Koh, Y.J., Lee, S.H., Lee, J., Kim, K.H., Kim, D., Koh, G.Y., and Yoo, O.J. (2013). Conditional ablation of LYVE-1+ cells unveils defensive roles of lymphatic vessels in intestine and lymph nodes. Blood 122, 2151–2161. 10.1182/BLOOD-2013-01-478941.

8. Piao, W., Xiong, Y., Li, L., Saxena, V., Smith, K.D., Hippen, K.L., Paluskievicz, C., Willsonshirkey, M., Blazar, B.R., Abdi, R., et al. (2020). Regulatory T Cells Condition Lymphatic Endothelia for Enhanced Transendothelial Migration. Cell Rep 30, 1052. 10.1016/J.CELREP.2019.12.083.

9. Saxena, V., Piao, W., Li, L., Paluskievicz, C., Xiong, Y., Simon, T., Lakhan, R., Brinkman, C.C., Walden, S., Hippen, K.L., et al. (2022). Treg tissue stability depends on lymphotoxin beta-receptor-and adenosine-receptor-driven lymphatic endothelial cell responses. Cell Rep 39, 110727. 10.1016/J.CELREP.2022.110727.

10. Cinti, I., and Denton, A.E. (2021). Lymphoid stromal cells-more than just a highway to humoral immunity. Oxf Open Immunol 2. 10.1093/OXFIMM/IQAB011.

11. Piao, W., Li, L., Paluskievicz, C., Hippen, K., Zhang, Y., Saxena, V., Shirkey, M.W., Blazar, B., Riella, L., and Bromberg, J. (2020). TREG USE THE PD-1-PD-L1 AXIS FOR LYMPHATIC TRANSENDOTHELIAL MIGRATION. Transplantation 104, S70–S70. 10.1097/01.TP.0000698604.45102.90.

12. Gousopoulos, E., Proulx, S.T., Bachmann, S.B., Scholl, J., Dionyssiou, D., Demiri, E., Halin, C., Dieterich, L.C., and Detmar, M. (2016). Regulatory T cell transfer ameliorates lymphedema and promotes lymphatic vessel function. JCI Insight 1. 10.1172/JCI.INSIGHT.89081.

13. Glasner, A., Rose, S.A., Sharma, R., Gudjonson, H., Chu, T., Green, J.A., Rampersaud, S., Valdez, I.K., Andretta, E.S., Dhillon, B.S., et al. (2023). Conserved transcriptional connectivity of regulatory T cells in the tumor microenvironment informs new combination cancer therapy strategies. Nat Immunol 24, 1020–1035. 10.1038/S41590-023-01504-2.

14. Ali, N., Zirak, B., Rodriguez, R.S., Pauli, M.L., Truong, H.A., Lai, K., Ahn, R., Corbin, K., Lowe, M.M., Scharschmidt, T.C., et al. (2017). Regulatory T Cells in Skin Facilitate Epithelial Stem Cell Differentiation. Cell 169, 1119–1129.e11. 10.1016/j.cell.2017.05.002.

15. Charbonnier, L.M., Wang, S., Georgiev, P., Sefik, E., and Chatila, T.A. (2015). Control of peripheral tolerance by regulatory T cell-intrinsic Notch signaling. Nat Immunol 16, 1162–1173. 10.1038/NI.3288.

16. Suchting, S., and Eichmann, A. (2009). Jagged Gives Endothelial Tip Cells an Edge. Cell 137, 988–990. 10.1016/J.CELL.2009.05.024.

17. Benedito, R., Roca, C., Sörensen, I., Adams, S., Gossler, A., Fruttiger, M., and Adams, R.H. (2009). The Notch Ligands Dll4 and Jagged1 Have Opposing Effects on Angiogenesis. Cell 137, 1124–1135. 10.1016/j.cell.2009.03.025.

18. Nakata, T., Shimizu, H., Nagata, S., Ito, G., Fujii, S., Suzuki, K., Kawamoto, A., Ishibashi, F., Kuno, R., Anzai, S., et al. (2017). Data showing proliferation and differentiation of intestinal epithelial cells under targeted depletion of Notch ligands in mouse intestine. Data Brief 10, 551–556. 10.1016/J.DIB.2016.12.045.

19. Souilhol, C., Li, X., Canham, L., Roddie, H., Pirri, D., Ayllon, B.T., Chambers, E. V, Dunning, M.J., Ariaans, M., Li, J., et al. (2020). JAG1-NOTCH4 Mechanosensing Drives Atherosclerosis. bioRxiv, 2020.05.15.097931. 10.1101/2020.05.15.097931.

20. Raya-Sandino, A., Lozada-Soto, K.M., Rajagopal, N., Garcia-Hernandez, V., Luissint, A.C., Brazil, J.C., Cui, G., Koval, M., Parkos, C.A., Nangia, S., et al. (2023). Claudin-23 reshapes epithelial tight junction architecture to regulate barrier function. Nat Commun 14. 10.1038/S41467-023-41999-9.

21. Cha, B., Geng, X., Mahamud, M.R., Zhang, J.Y., Chen, L., Kim, W., Jho, E. hoon, Kim, Y., Choi, D., Dixon, J.B., et al. (2018). Complementary Wnt Sources Regulate Lymphatic Vascular Development via PROX1-Dependent Wnt/β-Catenin Signaling. Cell Rep 25, 571–584.e5. 10.1016/J.CELREP.2018.09.049.

22. Lutze, G., Haarmann, A., Demanou Toukam, J.A., Buttler, K., Wilting, J., and Becker, J. (2019). Non-canonical WNT-signaling controls differentiation of lymphatics and extension lymphangiogenesis via RAC and JNK signaling. Sci Rep 9. 10.1038/S41598-019-41299-7.

23. Chai, Q., Onder, L., Scandella, E., Gil-Cruz, C., Perez-Shibayama, C., Cupovic, J., Danuser, R., Sparwasser, T., Luther, S.A., Thiel, V., et al. (2013). Maturation of Lymph Node Fibroblastic Reticular Cells from Myofibroblastic Precursors Is Critical for Antiviral Immunity. Immunity 38, 1013–1024. 10.1016/J.IMMUNI.2013.03.012.

24. Fletcher, A.L., Malhotra, D., Acton, S.E., Lukacs-Kornek, V., Bellemare-Pelletier, A., Curry, M., Armant, M., and Turley, S.J. (2011). Reproducible isolation of lymph node stromal cells reveals site-dependent differences in fibroblastic reticular cells. Front Immunol 2. 10.3389/FIMMU.2011.00035/ABSTRACT.

25. Pan, Y., and Xia, L. (2015). Emerging roles of podoplanin in vascular development and homeostasis. Front Med 9, 421–430. 10.1007/S11684-015-0424-9.

26. Maruyama, Y., Maruyama, K., Kato, Y., Kajiya, K., Moritoh, S., Yamamoto, K., Matsumoto, Y., Sawane, M., Kerjaschki, D., Nakazawa, T., et al. (2014). The Effect of Podoplanin Inhibition on Lymphangiogenesis Under Pathological Conditions. Invest Ophthalmol Vis Sci 55, 4813–4822. 10.1167/IOVS.13-13711.

27. Lucas, E.D., and Tamburini, B.A.J. (2019). Lymph node lymphatic endothelial cell expansion and contraction and the programming of the immune response. Front Immunol 10. 10.3389/FIMMU.2019.00036/PDF.

28. The optimum marker for the detection of lymphatic vessels (Review) https://www.spandidos-publications.com/10.3892/mco.2017.1356.

29. Olsson, Y., Svensjö, E., Arfors, K.E., and Hultström, D. (1975). Fluorescein labelled dextrans as tracers for vascular permeability studies in the nervous system. Acta Neuropathol 33, 45–50. 10.1007/BF00685963.

30. Liersch, R., Shin, J.W., Bayer, M., Schwöppe, C., Schliemann, C., Berdel, W.E., Mesters, R., and Detmar, M. (2012). Analysis of a novel highly metastatic melanoma cell line identifies osteopontin as a new lymphangiogenic factor. Int J Oncol 41, 1455–1463. 10.3892/IJO.2012.1548.

31. Lund, A.W., Wagner, M., Fankhauser, M., Steinskog, E.S., Broggi, M.A., Spranger, S., Gajewski, T.F., Alitalo, K., Eikesdal, H.P., Wiig, H., et al. (2016). Lymphatic vessels regulate immune microenvironments in human and murine melanoma. Journal of Clinical Investigation 126, 3389–3402. 10.1172/JCI79434.

32. Guo, R., Zhou, Q., Proulx, S.T., Wood, R., Ji, R.C., Ritchlin, C.T., Pytowski, B., Zhu, Z., Wang, Y.J., Schwarz, E.M., et al. (2009). Inhibition of lymphangiogenesis and lymphatic drainage via vascular endothelial growth factor receptor 3 blockade increases the severity of inflammation in a mouse model of chronic inflammatory arthritis. Arthritis Rheum 60, 2666–2676. 10.1002/ART.24764.

33. Ajithkumar, T., Parkinson, C., Fife, K., Corrie, P., and Jefferies, S. (2015). Evolving treatment options for melanoma brain metastases. Lancet Oncol 16, e486–e497. 10.1016/S1470-2045(15)00141-2.

34. Astarita, J.L., Dominguez, C.X., Tan, C., Guillen, J., Pauli, M.L., Labastida, R., Valle, J., Kleinschek, M., Lyons, J., and Zarrin, A.A. (2023). Treg specialization and functions beyond immune suppression. Clin Exp Immunol 211, 176. 10.1093/CEI/UXAC123.

35. Balint, K., Xiao, M., Pinnix, C.C., Soma, A., Veres, I., Juhasz, I., Brown, E.J., Capobianco, A.J., Herlyn, M., and Liu, Z.J. (2005). Activation of Notch1 signaling is required for β-catenin-mediated human primary melanoma progression. Journal of Clinical Investigation 115, 3166–3176. 10.1172/JCI25001.

36. Meng, J., Jiang, Y. zhou, Zhao, S., Tao, Y., Zhang, T., Wang, X., Zhang, Y., Sun, K., Yuan, M., Chen, J., et al. (2022). Tumor-derived Jagged1 promotes cancer progression through immune evasion. Cell Rep 38, 110492. 10.1016/J.CELREP.2022.110492.

37. Qiao, X., Ma, B., Sun, W., Zhang, N., Liu, Y., Jia, L., and Liu, C. (2022). JAG1 is associated with the prognosis and metastasis in breast cancer. Sci Rep 12. 10.1038/S41598-022-26241-8.

38. Gordon, B., Swaminathan, B., Naiche, L., and Kitajewski, J. (2023). Abstract P6-14-10: Jagged-1 promotes breast cancer metastasis through the lymphatic system. Cancer Res 83, P6–14–10. 10.1158/1538-7445.SABCS22-P6-14-10.

39. Gillot, L., Baudin, L., Rouaud, L., Kridelka, F., and Noël, A. (2021). The pre-metastatic niche in lymph nodes: formation and characteristics. Cell Mol Life Sci 78, 5987–6002. 10.1007/S00018-021-03873-Z.

40. Herrera, J.L., and Komatsu, M. (2024). Akt3 activation by R-Ras in an endothelial cell enforces quiescence and barrier stability of neighboring endothelial cells via Jagged1. Cell Rep 43, 113837. 10.1016/J.CELREP.2024.113837.

41. Fatima, A., Culver, A., Culver, F., Liu, T., Dietz, W.H., Thomson, B.R., Hadjantonakis, A.K., Quaggin, S.E., and Kume, T. (2014). Murine Notch1 is required for lymphatic vascular morphogenesis during development. Developmental Dynamics 243, 957–964. 10.1002/DVDY.24129.

42. Rantakari, P., Auvinen, K., Jäppinen, N., Kapraali, M., Valtonen, J., Karikoski, M., Gerke, H., Iftakhar-E-Khuda, I., Keuschnigg, J., Umemoto, E., et al. (2015). The endothelial protein PLVAP in lymphatics controls the entry of lymphocytes and antigens into lymph nodes. Nat Immunol 16, 386–396. 10.1038/NI.3101.

43. Weigel, C., Bellaci, J., and Spiegel, S. (2023). Sphingosine-1-phosphate and its receptors in vascular endothelial and lymphatic barrier function. J Biol Chem 299, 104775. 10.1016/J.JBC.2023.104775.

44. Aho, S. (2004). Soluble form of Jagged1: Unique product of epithelial keratinocytes and a regulator of keratinocyte differentiation. J Cell Biochem 92, 1271–1281. 10.1002/JCB.20125.

45. Nanaware, P.P., Khan, Z.N., Clement, C.C., Shetty, M., Mota, I., Seltzer, E.S., Dzieciatkowska, M., Gamboni, F., D’Alessandro, A., Ng, C., et al. (2024). Role of the afferent lymph as an immunological conduit to analyze tissue antigenic and inflammatory load. Cell Rep 43, 114311. 10.1016/J.CELREP.2024.114311/ATTACHMENT/8070E23F-DC65-4C58-8516-F8F5505ADEA6/MMC1.

46. Flier, J.S., Underhill, L.H., and Dvorak, H.F. (1986). Tumors: wounds that do not heal. Similarities between tumor stroma generation and wound healing. N Engl J Med 315, 1650–1659. 10.1056/NEJM198612253152606.

47. Cohen, J.N., Gouirand, V., Macon, C.E., Lowe, M.M., Boothby, I.C., Moreau, J.M., Gratz, I.K., Stoecklinger, A., Weaver, C.T., Sharpe, A.H., et al. (2024). Regulatory T cells in skin mediate immune privilege of the hair follicle stem cell niche. Sci Immunol 9. 10.1126/sciimmunol.adh0152.

48. Bankhead, P., Loughrey, M.B., Fernández, J.A., Dombrowski, Y., McArt, D.G., Dunne, P.D., McQuaid, S., Gray, R.T., Murray, L.J., Coleman, H.G., et al. (2017). QuPath: Open source software for digital pathology image analysis. Sci Rep 7. 10.1038/S41598-017-17204-5.

49. Bolger, A.M., Lohse, M., and Usadel, B. (2014). Trimmomatic: a flexible trimmer for Illumina sequence data. Bioinformatics 30, 2114–2120. 10.1093/BIOINFORMATICS/BTU170.

50. Dobin, A., Davis, C.A., Schlesinger, F., Drenkow, J., Zaleski, C., Jha, S., Batut, P., Chaisson, M., and Gingeras, T.R. (2013). STAR: ultrafast universal RNA-seq aligner. Bioinformatics 29, 15–21. 10.1093/BIOINFORMATICS/BTS635.

51. Liao, Y., Smyth, G.K., and Shi, W. (2013). The Subread aligner: fast, accurate and scalable read mapping by seed-and-vote. Nucleic Acids Res 41. 10.1093/NAR/GKT214.

52. GitHub - kevinblighe/EnhancedVolcano: Publication-ready volcano plots with enhanced colouring and labeling https://github.com/kevinblighe/EnhancedVolcano.

53. Chen, E.Y., Tan, C.M., Kou, Y., Duan, Q., Wang, Z., Meirelles, G. V., Clark, N.R., and Ma’ayan, A. (2013). Enrichr: interactive and collaborative HTML5 gene list enrichment analysis tool. BMC Bioinformatics 14. 10.1186/1471-2105-14-128.

54. CRAN - Package pheatmap https://cran.r-project.org/web/packages/pheatmap/index.html.

55. Smedley, D., Haider, S., Durinck, S., Pandini, L., Provero, P., Allen, J., Arnaiz, O., Awedh, M.H., Baldock, R., Barbiera, G., et al. (2015). The BioMart community portal: an innovative alternative to large, centralized data repositories. Nucleic Acids Res 43, W589–W598. 10.1093/NAR/GKV350.

56. Foroutan, M., Bhuva, D.D., Lyu, R., Horan, K., Cursons, J., and Davis, M.J. (2018). Single sample scoring of molecular phenotypes. BMC Bioinformatics 19. 10.1186/S12859-018-2435-4.

57. Therneau, T.M. (2024). Survival Analysis [R package survival version 3.7–0].

58. Drawing Survival Curves using “ggplot2” [R package survminer version 0.4.9] (2021).

59. Create Elegant Data Visualisations Using the Grammar of Graphics • ggplot2 https://ggplot2.tidyverse.org/.

